# *MSH1* is required for maintenance of the low mutation rates in plant mitochondrial and plastid genomes

**DOI:** 10.1101/2020.02.13.947598

**Authors:** Zhiqiang Wu, Gus Waneka, Amanda K. Broz, Connor R. King, Daniel B. Sloan

**Affiliations:** Shenzhen Branch, Guangdong Laboratory for Lingnan Modern Agriculture, Genome Analysis Laboratory of the Ministry of Agriculture, Agricultural Genomics Institute at Shenzhen, Chinese Academy of Agricultural Sciences, Shenzhen, China 518120; Department of Biology, Colorado State University, Fort Collins, CO 80523

## Abstract

Mitochondrial and plastid genomes in land plants exhibit some of the slowest rates of sequence evolution observed in any eukaryotic genome, suggesting an exceptional ability to prevent or correct mutations. However, the mechanisms responsible for this extreme fidelity remain unclear. We tested seven candidate genes involved in cytoplasmic DNA replication, recombination, and repair (*POLIA*, *POLIB*, *MSH1*, *RECA3*, *UNG*, *FPG*, and *OGG1*) for effects on mutation rates in the model angiosperm *Arabidopsis thaliana* by applying a highly accurate DNA sequencing technique (duplex sequencing) that can detect newly arisen mitochondrial and plastid mutations still at low heteroplasmic frequencies. We find that disrupting *MSH1* (but not the other candidate genes) leads to massive increases in the frequency of point mutations and small indels and changes to the mutation spectrum in mitochondrial and plastid DNA. We also used droplet digital PCR to show transmission of *de novo* heteroplasmies across generations in *msh1* mutants, confirming a contribution to heritable mutation rates. This dual-targeted gene is part of an enigmatic lineage within the *mutS* mismatch repair family that we find is also present outside of green plants in multiple eukaryotic groups (stramenopiles, alveolates, haptophytes, and cryptomonads), as well as certain bacteria and viruses. *MSH1* has previously been shown to limit ectopic recombination in plant cytoplasmic genomes. Our results point to a broader role in recognition and correction of errors in plant mitochondrial and plastid DNA sequence, leading to greatly suppressed mutation rates perhaps via initiation of double-stranded breaks and repair pathways based on faithful homologous recombination.

## INTRODUCTION

It has been apparent for more than 30 years that rates of nucleotide substitution in plant mitochondrial and plastid genomes are unusually low (1, 2). In angiosperms, mitochondrial and plastid genomes have synonymous substitution rates that are on average approximately 16-fold and 5-fold slower than the nucleus, respectively (3). The fact that these low rates are evident even at sites that are subject to relatively small amounts of purifying selection (4, 5) suggests that they are the result of very low underlying mutation rates – a surprising observation especially when contrasted with the rapid accumulation of mitochondrial mutations in many eukaryotic lineages (6, 7).

Although the genetic mechanisms that enable plants to achieve such faithful replication and transmission of cytoplasmic DNA sequences have not been determined, a number of hypotheses can be envisioned. One possibility is that the DNA polymerases responsible for replicating mitochondrial and plastid DNA (8) might have unusually high fidelity. However, *in vitro* assays with the two partially redundant bacterial-like organellar DNA polymerases in *Arabidopsis thaliana*, PolIA (At1g50840) and PolIB (At3g20540), have indicated that they are highly error-prone (9), with misincorporation rates that exceed those of Pol γ, the enzyme responsible for replicating DNA in the rapidly mutating mitochondrial genomes of humans (10). PolIB has a measured error rate (5.45 × 10^−4^ per bp) that is 7.5-fold higher than that of PolIA (7.26 × 10^−5^ per bp). Therefore, although knocking out both of these genes results in lethality (8), disrupting one of the two polymerases to make the cell rely on the other may provide an opportunity to investigate the effects of polymerase misincorporation on the overall mutation rate.

It is also possible that plant mitochondria and plastids are unusually effective at preventing or repairing DNA damage resulting from common mechanisms such as guanine oxidation (e.g., 8-oxo-G) and cytosine deamination (uracil). Like most organisms, plants encode dedicated enzymes to recognize these forms of damage and initiate base-excision repair. In *Arabidopsis*, a pair of enzymes (FPG [At1g52500] and OGG1 [At1g21710]) both appear to function in repair of 8-oxo-G in mitochondrial and nuclear DNA, as evidenced by increased oxidative damage in the double mutant background (11), and uracil N-glycosylase (UNG [At3g18630]) recognizes and removes uracil in all three genomic compartments (12, 13). A recent study investigated the effects of knocking out *UNG* on mitochondrial sequence variation in *Arabidopsis* but did not find any nucleotide substitutions that rose to high frequency nor any difference in variant frequencies relative to wild type in 10-generation mutation accumulation lines (14). The apparent low fidelity of plant organellar DNA polymerases and tolerance of disruptions to the *UNG* base-excision repair pathway suggest that other mechanisms are at play in dealing with mismatches and DNA damage.

Mitochondrial and plastid genomes are present in numerous copies per cell, and it is often hypothesized that recombination and homology-directed repair (HDR) may eliminate mutations and damaged bases in plant cytoplasmic genomes (9, 14–17). In both mitochondria and plastids, there is extensive recombination and gene conversion between homologous DNA sequences (18–20), and the large inverted repeats in plastid genomes have slower sequence evolution than single-copy regions (1, 21), suggesting that increased availability of homologous templates may improve the accuracy of error correction. The extensive recombinational dynamics in plant mitochondrial genomes often extend to short repeat sequences, resulting in structural rearrangements. As such, the slow rate of sequence evolution in these genomes is juxtaposed with rapid structural change (20).

*MutS Homolog 1* (*MSH1* [At3g24320]) is involved in recombination in both mitochondria and plastids and represents a natural candidate for maintaining low mutation rates in plant cytoplasmic genomes. Plants homozygous for mutated copies of this gene often develop variegated leaf phenotypes that subsequently follow a pattern of maternal inheritance, indicating alterations in cytoplasmic genomes (22–25). *MSH1* is distinguished from other members of the larger *mutS* mismatch repair (MMR) gene family by an unusual C-terminal domain predicted to be a GIY-YIG endonuclease (26). This observation led Christensen (16) to hypothesize that MSH1 recognizes mismatches or DNA damage and introduces double-stranded breaks (DSBs) at those sites as a means to initiate accurate repair via HDR. However, analysis and sequencing of mitochondrial and plastid genomes in *msh1* mutants has not detected base substitutions or small indels (24, 25, 27). Instead, characterization of cytoplasmic genomes in these mutants has revealed structural rearrangements resulting from ectopic recombination between small repeats (24, 25, 28). These findings have led to the prevailing view that the primary role of MSH1 is in recombination surveillance rather than in correction of mismatches or damaged bases (24, 26, 29–32), as is the case for some other members of the *mutS* gene family (33). Numerous other genes have also been identified as playing a role in mitochondrial and plastid recombination (20). One example is the mitochondrial-targeted *RECA3* gene (At3g10140), with *recA3* and *msh1* mutants exhibiting similar but non-identical effects in terms of repeat-mediated rearrangements and aberrant growth phenotypes (29, 34).

It is striking that so many genes have been identified in controlling the structural stability of plant mitochondrial and plastid genomes (20) and yet researchers have not been able to identify any gene knockouts in plants that lead to increased cytoplasmic mutation rates despite the many promising hypotheses and candidates. This gap may reflect the inherent challenges in studying rare mutational events in long-lived multicellular organisms. The advent of high-throughput DNA sequencing has raised the possibility of using deep sequencing coverage to catch *de novo* mutations essentially as they arise and are still present at extremely low frequencies among the many cytoplasmic genome copies that are found within cells and tissue samples (heteroplasmy). However, the error rate of standard sequencing technologies such as Illumina is relatively high – often above 10^−3^ errors per bp and much worse in certain sequence contexts (35) – setting a problematic noise threshold for accurate detection of rare variants. Fortunately, numerous specialized methods have been introduced to improve these error rates (36). The most accurate technique is known as duplex sequencing (37), which entails tagging both ends of each original DNA fragment with adapters containing random barcodes such that it is possible to obtain a consensus from multiple reads originating from the same biological molecule, including those from each of the two complementary DNA strands. Duplex sequencing has been found to reduce error rates approximately 10,000-fold to levels below 10^−7^ errors per bp (37), opening the door for accurate detection of extremely rare variants.

Here, we have applied duplex sequencing to detect *de novo* mitochondrial and plastid mutations in wild type *Arabidopsis* and a number of mutant backgrounds carrying disrupted copies of key nuclear candidate genes involved in cytoplasmic DNA recombination, replication, and repair (RRR). We find that, of these candidates, only *msh1* mutants show massive increases in rates of point mutations and small indels in cytoplasmic genomes, identifying this gene as a key player in maintaining the remarkably low mutation rates in plant mitochondria and plastids.

## RESULTS

### Detection of mitochondrial and plastid mutations in wild type *Arabidopsis*

We modified the standard duplex sequencing protocol to include treatment with repair enzymes that correct single-stranded DNA damage and established a noise threshold of approximately 2 × 10^−8^ sequencing errors per bp using *E. coli* samples derived from single colonies (Supplementary Text, Tables S1 and S2). We then applied this sequencing method to purified mitochondrial and plastid DNA from *Arabidopsis thaliana* Col-0 rosette tissue to provide a baseline characterization of the variant spectrum in wild type plants. Following the removal of spurious variants that resulted from contaminating nuclear copies of mitochondrial- or plastid-derived sequences (known as NUMTs and NUPTs (38), respectively) or from chimeric molecules produced by recombination between non-identical repeats, we found that almost all single-nucleotide variant (SNV) types were present at a frequency of less than 10^−7^, suggesting that levels of standing variation were generally at or near the noise threshold despite the extreme sensitivity of this method. The one obvious exception was GC→AT transitions in mtDNA, which were detected at a mean frequency of 3.8 × 10^−7^ across three biological replicates. The dominance of GC→AT transitions in the mitochondrial mutation spectrum was further supported by subsequent sequencing of 24 additional wild type control replicates (Fig. 1) that were part of later experiments investigating individual candidate genes. GC→AT transitions also tended to be the most common type of SNV in plastid DNA samples but at a level that was almost an order of magnitude lower than that observed in mitochondrial samples (mean frequency of 4.6 × 10^−8^).

**Figure 1.**
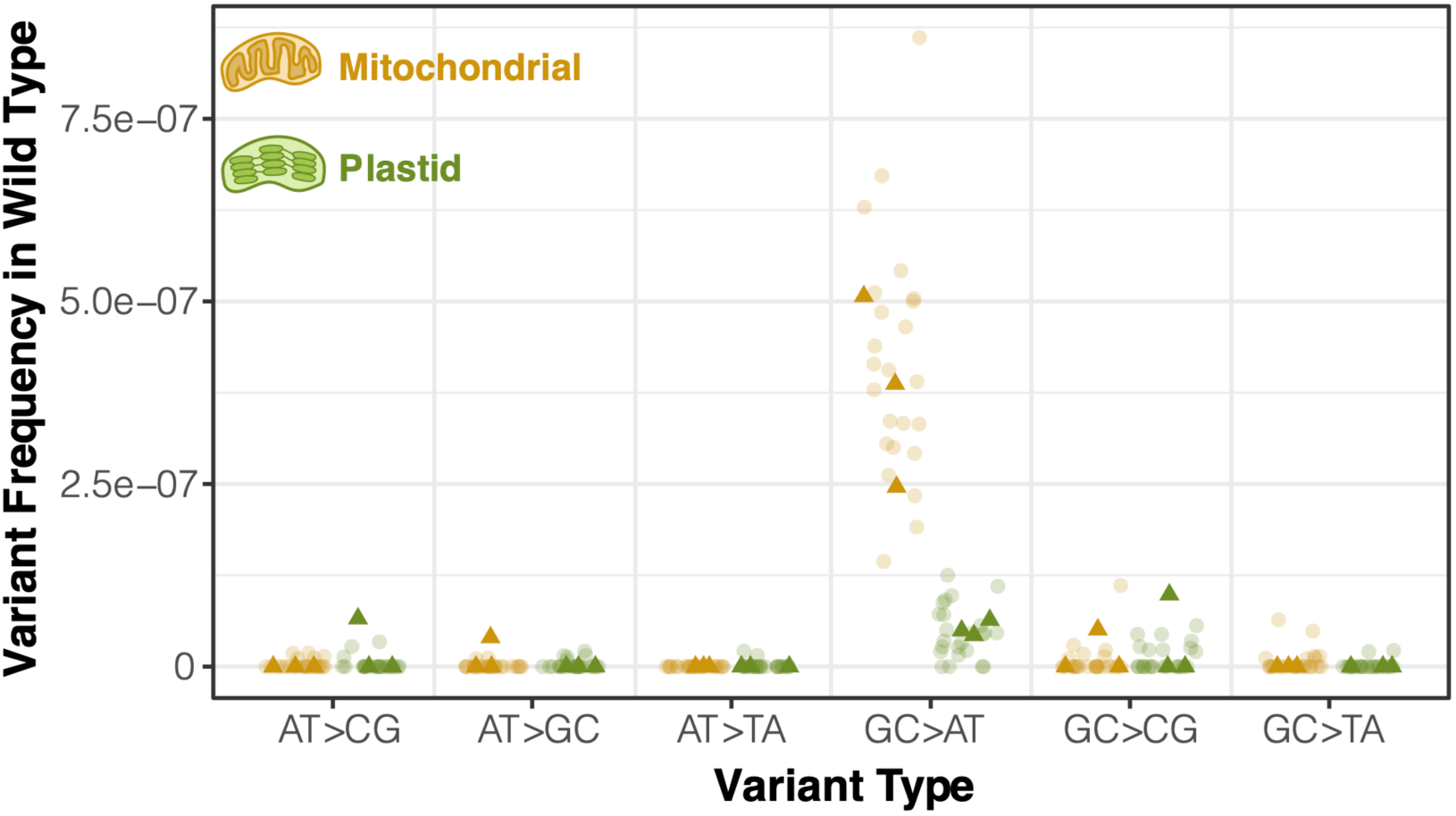
Observed frequency of mitochondrial and plastid SNVs in wild-type *Arabidopsis* tissue based on duplex sequencing. Dark triangles represent three biological replicate families of wild-type *A. thaliana* Col-0. Lighter circles are F3 families derived from homozygous wild-type plants that segregated out from a heterozygous parent containing one mutant copy of an RRR candidate gene. Variant frequencies are calculated as the total number of observed mismatches in mapped duplex consensus sequences divided by the total bp of sequence coverage.

### Screen of candidate genes reveals greatly increased frequency of mitochondrial and plastid mutations in *msh1* knockout

To test the effects of disrupting key RRR genes (Table S3) on cytoplasmic mutation rates, we applied crossing designs that enabled direct comparisons between families of homozygous mutants and matched wild type controls that all inherited their cytoplasmic genomes from the same grandparent (Fig. 2). We performed duplex sequencing with purified mitochondrial and plastid DNA from the resulting samples, generating a total of 1.2 Tbp of raw Illumina reads that were collapsed down into 10 Gbp of processed and mapped duplex consensus sequence (DCS) data (Table S4). Many of the candidate genes have been previously found to affect structural stability in the mitochondrial genome (8, 29, 34). Consistent with these expected structural effects, we found that *msh1*, *recA3*, and *polIb* mutants all showed their own distinct patterns of large and repeatable shifts in coverage in regions of the mitochondrial genome (Fig. 3). Shifts in coverage were weaker in *polIa* mutants and not detected in *ung* mutants or *fpg/ogg1* double mutants. Such coverage variation was not found in the plastid genome for any of the mutants (Fig. S1).

**Figure 2.**
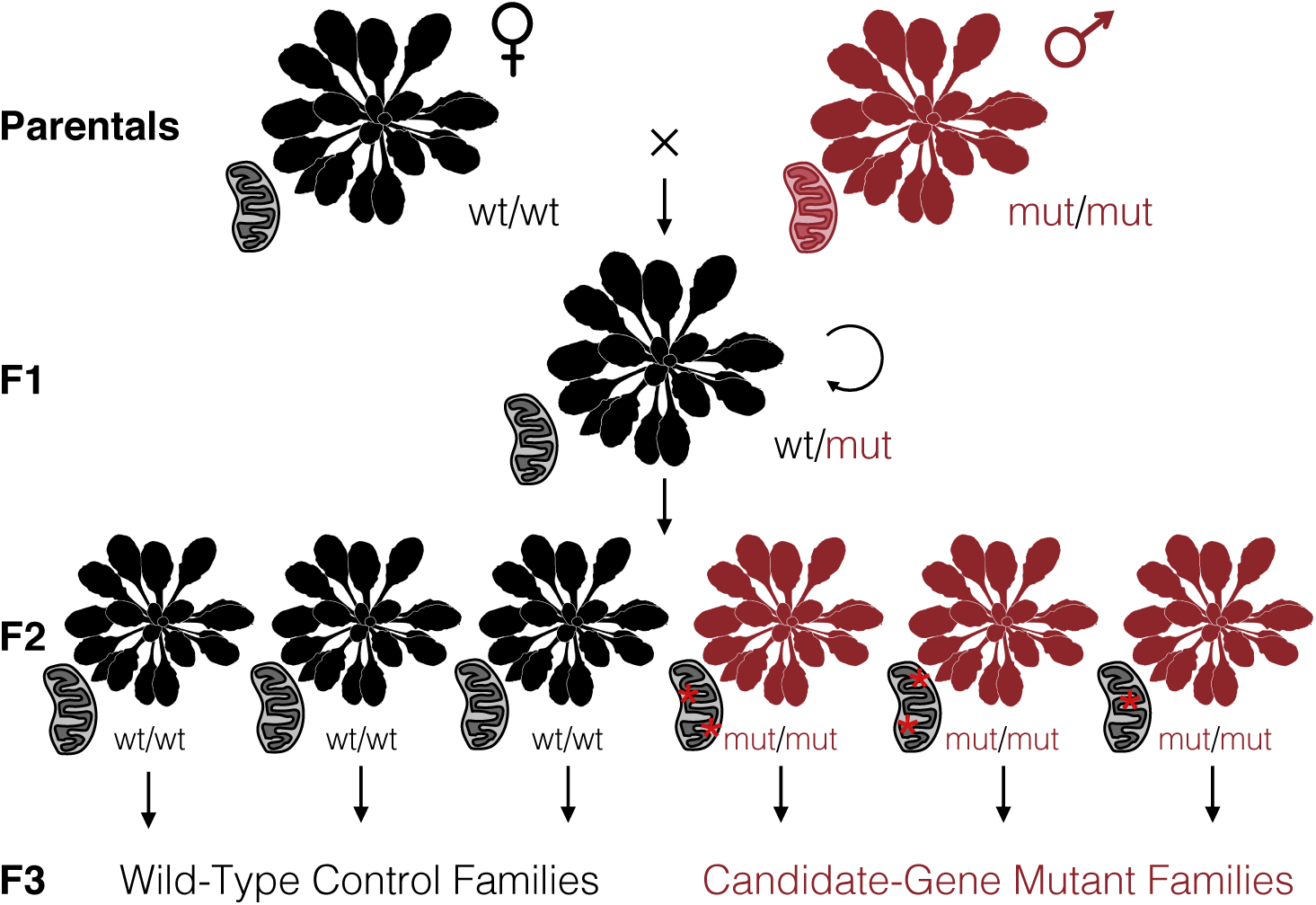
Crossing design to test candidate nuclear genes involved in RRR of cytoplasmic genomes. Using a wild-type maternal plant (black) and either a homozygous mutant (red) or heterozygous pollen donor, we generated a heterozygous F1 individual that carried cytoplasmic genomes inherited from a wild-type lineage (as indicated by the black mitochondrion). After selfing the F1, we genotyped the resulting F2 progeny to identify three homozygous mutants and three homozygous wild-type individuals. Given that the mutations in candidate RRR genes are expected to be recessive, the F2 generation would be the first in which the sampled cytoplasmic genomes were exposed to the effects (red asterisks) of these mutants. The identified F2 individuals were each allowed to self-fertilize and set seed to produce multiple F3 families that all inherited their cytoplasmic genomes from the same F1 grandparent. The F3 families were used for purification of mitochondrial and plastid DNA for duplex sequencing. Sequencing was performed on three replicate families for each genotype. *Arabidopsis* silhouette image is from PhyloPic (Mason McNair).

**Figure 3.**
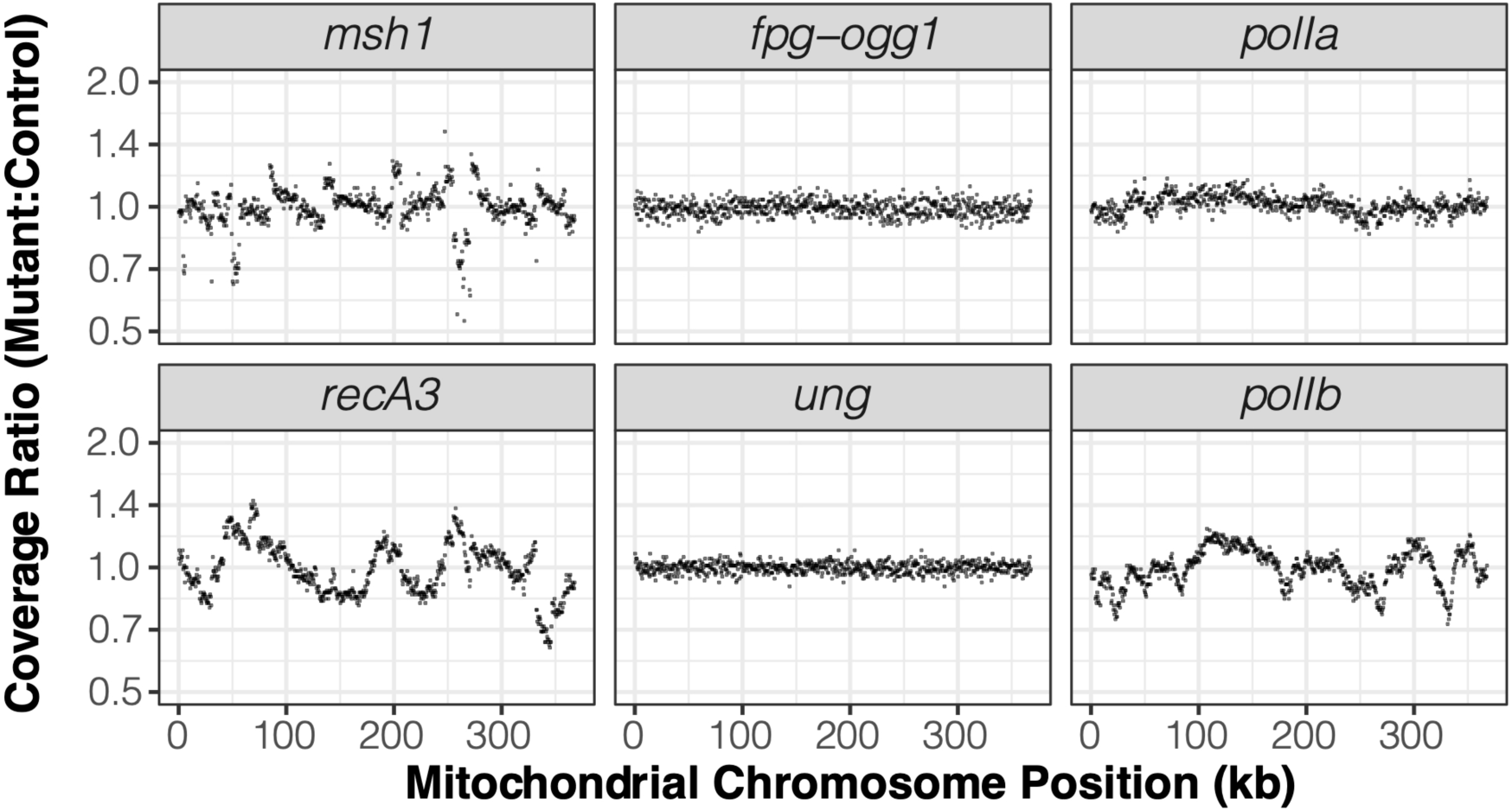
Sequencing coverage variation across the mitochondrial genome in mutants relative to their matched wild type controls. Each panel represents an average of three biological replicates. The reported ratios are based on counts per million mapped reads in 500-bp windows. The *msh1* mutant line reported in this figure is CS3246.

Most of the analyzed candidate genes did not have detectable effects on cytoplasmic mutation rates. Despite the difference in measured misincorporation rates for PolIA and PolIB *in vitro* (9), we did not find that disrupting either of these genes had an effect on the frequencies of SNVs or small indels *in vivo* (Figs. 4 and S2). Likewise, *ung* mutants and *fpg/ogg1* double mutants did not exhibit any detectable increase in sequence variants. In *recA3* mutants, there was a weak trend towards higher rates of mitochondrial SNVs and small indels compared to wild type controls (Fig. S2), but neither of these effects were statistically significant.

**Figure 4.**
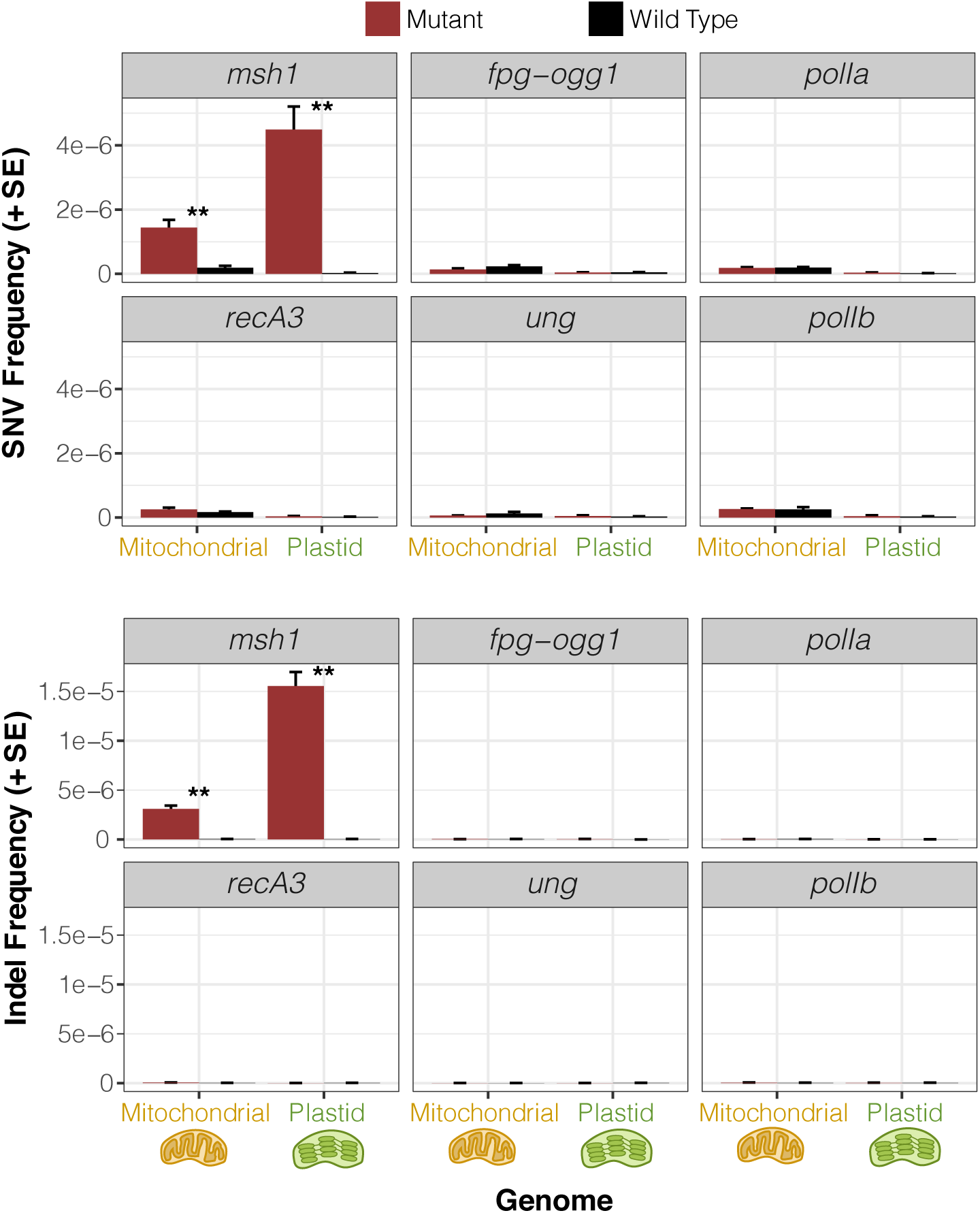
Observed frequency of mitochondrial and plastid SNVs (top) and indels (bottom) based on duplex sequencing in *Arabidopsis* mutant backgrounds for various RRR genes compared to matched wild type controls. Variant frequencies are calculated as the total number of observed mismatches or indels in mapped duplex consensus sequences divided by the total bp of sequence coverage. Means and standard errors are based on three replicate F3 families for each genotype (see Fig. 2). The *msh1* mutant line reported in this figure is CS3246. Significant differences between mutant and wild type genotypes at a level of *P* < 0.01 (*t*-tests on log-transformed values) are indicated by **. All other comparisons were non-significant (*P* > 0.05).

Unlike the rest of the candidate genes, the *msh1* mutant line (CS3246) exhibited a striking increase in SNVs compared to wild type controls – close to 10-fold in the mitochondrial genome and more than 100-fold in the plastid genome (Figs. 4 and S2). The *msh1* mutation spectrum in both mitochondrial and plastid DNA was dominated by transitions. GC→AT substitutions remained the most common mitochondrial SNV in *msh1* mutants, but there was a disproportionate increase in AT→GC variants such that both transition types reached comparable levels (Fig. 5). The increased frequency of AT→GC transitions in *msh1* mutants was even more dramatic for plastid DNA, making them by far the most abundant type of SNV. Disruption of *MSH1* also affected transversion rates, with substantial increases in GC→TA and AT→CG SNVs in both genomes (Fig. 5).

**Figure 5.**
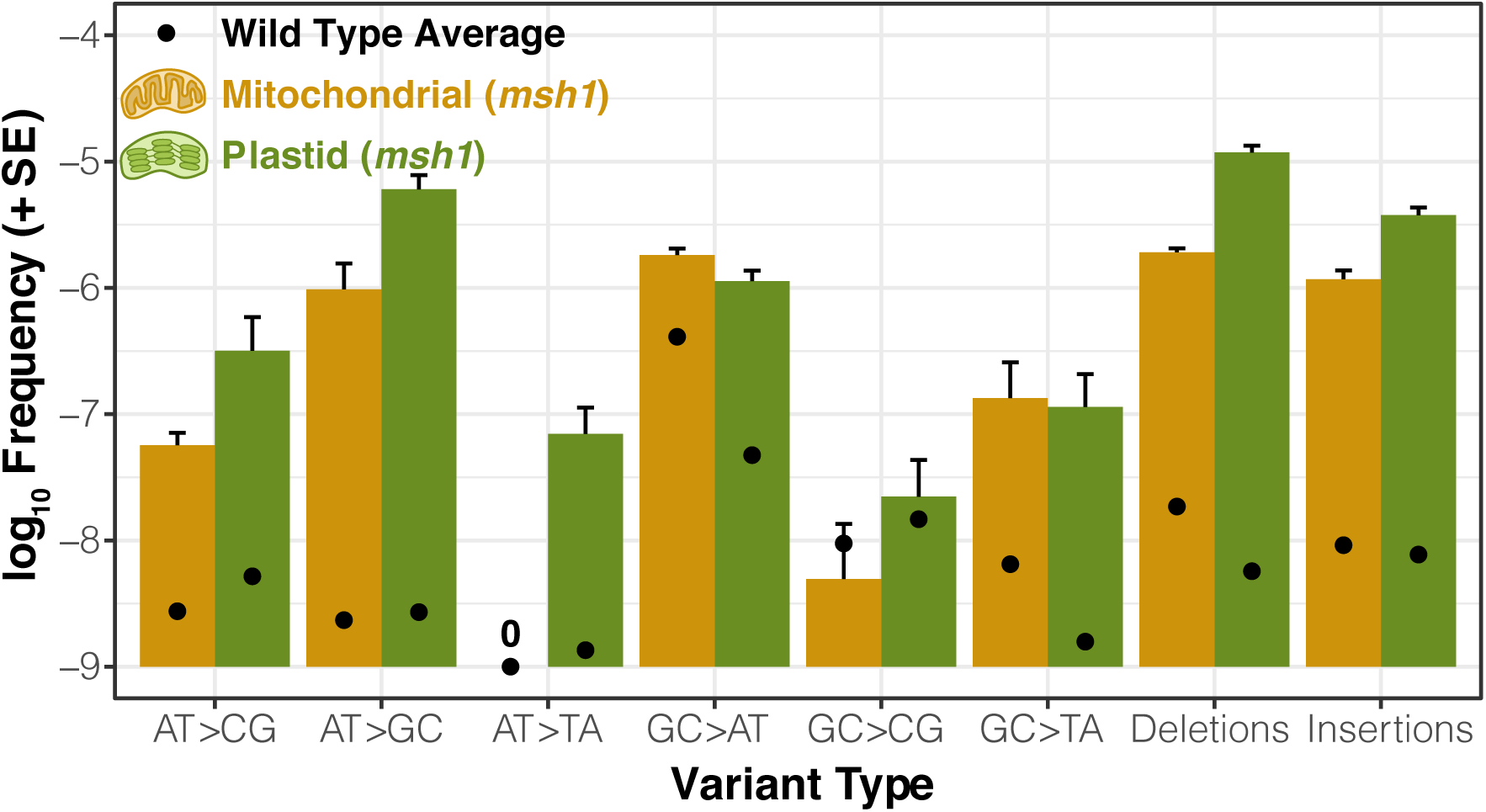
Spectrum of point mutations and indels in *Arabidopsis msh1* mutants (CS3246 allele). Variant frequencies are calculated as the total number of observed mismatches or indels in mapped duplex consensus sequences divided by the total bp of sequence coverage. Means and standard errors are based on three replicate F3 families. Black points represent the mean variant frequency calculated from all wild type libraries (Fig. 1). Note that the “0” indicates that no mitochondrial AT→TA transversions were observed in any of the *msh1*-CS3246 or wild type libraries.

The rate of small indel mutations also increased dramatically in the *msh1* mutant line, with indel frequencies jumping approximately two or three orders of magnitude in the mitochondrial and plastid genomes, respectively (Figs. 4 and S2). The indels in *msh1* mutants overwhelmingly occurred in homopolymer regions (i.e., single-nucleotide repeats). On average, 90.8% of observed indels from mapped DCS data in the mitochondrial genome and 97.7% in the plastid genome were in homopolymers at least 6 bp in length. There was a clear bias towards deletions in the *msh1* mutants, with deletions 1.6-fold and 3.1-fold more abundant than insertions on average in mitochondrial and plastid genomes, respectively (Fig. 5).

### Confirmation of *msh1* mutator effects in additional mutant backgrounds

To verify that disruption of *msh1* was indeed responsible for the observed elevation in mitochondrial and plastid mutation rates, we repeated our crossing design and duplex sequencing analysis with two additional independently derived mutant alleles in this gene (Table S3). All three *msh1* mutant backgrounds showed the same qualitative pattern of increased SNVs, small indels, and structural variation (Figs. S3 and S4). The magnitude of these effects was equivalent in the initial mutant line (CS3246) and a second mutant (CS3372), which both harbor point mutations that appear to generate null *msh1* alleles (24). In contrast, the increases in sequence and structural variation were much smaller for a third *msh1* allele (SALK_046763). The SALK_046763 mutants also exhibited weaker phenotypic effects, with lower rates of visible leaf variegation (Fig. S5). The 3357-bp *MSH1* coding sequence is distributed across 22 exons, and this mutant allele carries a T-DNA insertion in the eighth intron (24), which we reasoned might reduce but not eliminate expression of functional MSH1 protein, resulting in weaker effects on phenotype and mutation rates. In support of this prediction, cDNA sequencing across the boundary between exons 8 and 9 confirmed the presence of properly spliced transcripts in homozygous SALK_046763 mutants despite the large T-DNA insertion in the intron (Fig. S6a. Furthermore, quantitative reverse-transcriptase PCR (qRT-PCR) showed that expression levels in leaf tissue were roughly 5-fold lower than in wild type individuals (Fig. S6b). Therefore, reducing the expression level of *MSH1* also appears to increase cytoplasmic mutation rates, though to a lesser degree than effective knockouts.

### Inheritance of *msh1*-induced heteroplasmies

Because we performed our duplex sequencing analysis on whole-rosette tissue, it was not immediately clear whether the increase in observed mutations included changes that could be transmitted across generations or only variants that accumulated in vegetative tissue and would not be inherited. The majority of SNVs (~80%) in *msh1* mutants and all SNVs in their matched wild type controls were detected in only a single DCS read family (Dataset S1), implying very low heteroplasmic frequency in the pooled F3 tissue as would be expected for new mutations. However, we did identify a total of 433 SNVs across the *msh1* mutant samples that were each supported by multiple DCS read families (i.e., distinct biological molecules in the original DNA samples), in some cases reaching frequencies of >2%. We reasoned that to be found at such frequencies in a pool of tissue from dozens of F3 individuals, a variant likely had to have occurred in the F2 parent and been inherited in a heteroplasmic state by multiple F3s. Although the individuals used for duplex sequencing were sacrificed in the process of extracting mitochondrial and plastid DNA from whole rosettes, we had collected F4 seed from siblings of the F3 plants that were grown up in parallel. Therefore, we developed droplet digital PCR (ddPCR) markers to test for the inheritance of some identified high-frequency SNVs in the F4 generation.

We assayed five SNVs with ddPCR markers, each of which was found at substantial frequencies in the corresponding F3 cytoplasmic DNA sample (1.4 to 14.3%), confirming the variant identification from our duplex sequencing. As controls, we sampled F4 *msh1* mutants derived from other F3 families that did not show evidence of the variant in duplex sequencing data. All of these controls exhibited a frequency of below 0.2%, which we considered the noise threshold for the assay. For two of the five markers (one mitochondrial and one plastid), we were also able to detect the SNV in DNA samples from individual F4 plants. The frequency of these heteroplasmic mutations varied dramatically across F4 individuals – anywhere from below the noise threshold to as high as apparent homoplasmy (>99.9%) in one case (Fig. 6). These high frequencies indicate the potential for *de novo* mutations to spread to majority status remarkably fast, and they represent clear evidence that *de novo* cytoplasmic mutations can occur in meristematic tissue in an *msh1* background and be transmitted across generations, thereby increasing the heritable mutation rate. For the other three SNVs, we did not detect the heteroplasmic mutation in a sample of eight F4 individuals, which could indicate that the variant was restricted to vegetative tissue in a single individual within the F3 pool. However, we suspect that it is more likely that the negative F4 individuals lost the corresponding variant via a heteroplasmic sorting process or descended from a subset of F3 parents that did not carry it.

**Figure 6.**
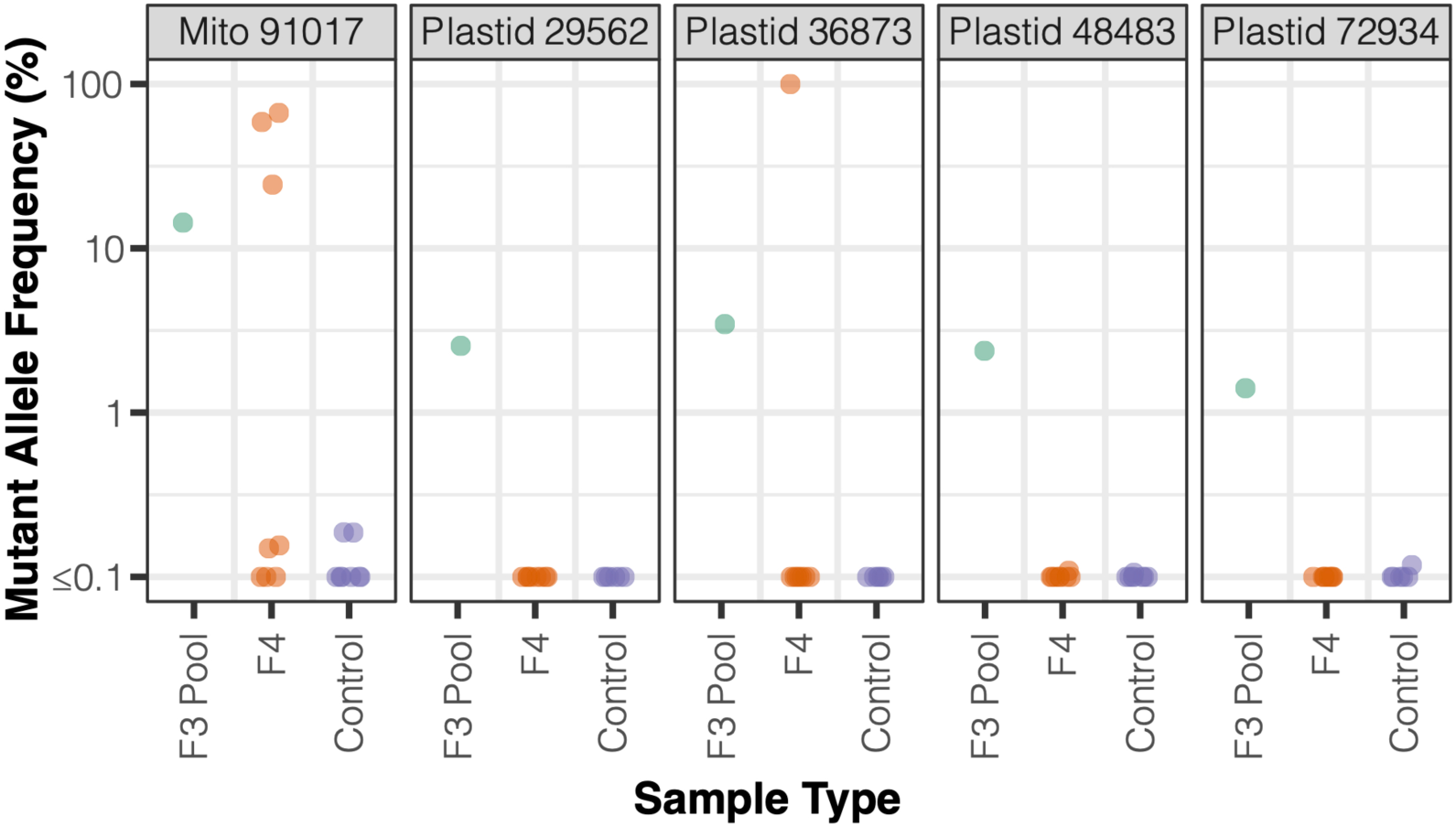
Estimates of heteroplasmic frequency of select SNVs using ddPCR. The F3 pool is the same *msh1*-CS3246 mutant mitochondrial or plastid DNA sample in which the SNV was discovered by duplex sequencing. F4 indicates plants descended from the F3 family used in duplex sequencing. Controls are F4 plants from *msh1*-CS3246 mutant lines other than the one in which the SNV was originally discovered.

### Plant *MSH1* is a part of widely distributed gene family in diverse eukaryotic lineages, as well some bacteria and viruses

*MSH1* is divergent in sequence and domain architecture relative to all other members of the *mutS* MMR gene family (39). Although named after the *MSH1* gene in yeast, which also functions in mitochondrial DNA repair (40), plant *MSH1* is from an entirely different part of the large *mutS* family (39). It is known to be widely present across green plants (41), but its evolutionary history beyond that is unclear. Taxon-specific searches of public genomic and metagenomic repositories failed to detect copies of *MSH1* in red algae and glaucophytes, the other two major lineages of Archaeplastida. Likewise, we did not find evidence of this gene in Amorphea (which includes Amoebozoa, animals, fungi, and related protists). Although these initial results implied a distribution that might be truly restricted to green plants (Viridiplantae), searches of other major eukaryotic lineages found that plant-like *MSH1* homologs carrying the characteristic GIY-YIG endonuclease domain were present in numerous groups – specifically stramenopiles, alveolates, haptophytes, and cryptomonads (Fig. 7). More surprisingly, we found that it was present in the genomes of two closely related bacterial species within the Cellvibrionaceae (Gammaproteobacteria) and another gammaproteobacterium of uncertain classification, as well as some unclassified viruses curated from environmental and metagenomic datasets (42, 43). Phylogenetic analysis confirmed that these sequences represented a well-resolved clade within the *mutS* family (Fig. 7). Therefore, the plant-like *MSH1* gene appears to have an unusually disjunct distribution across diverse lineages in the tree of life.

**Figure 7.**
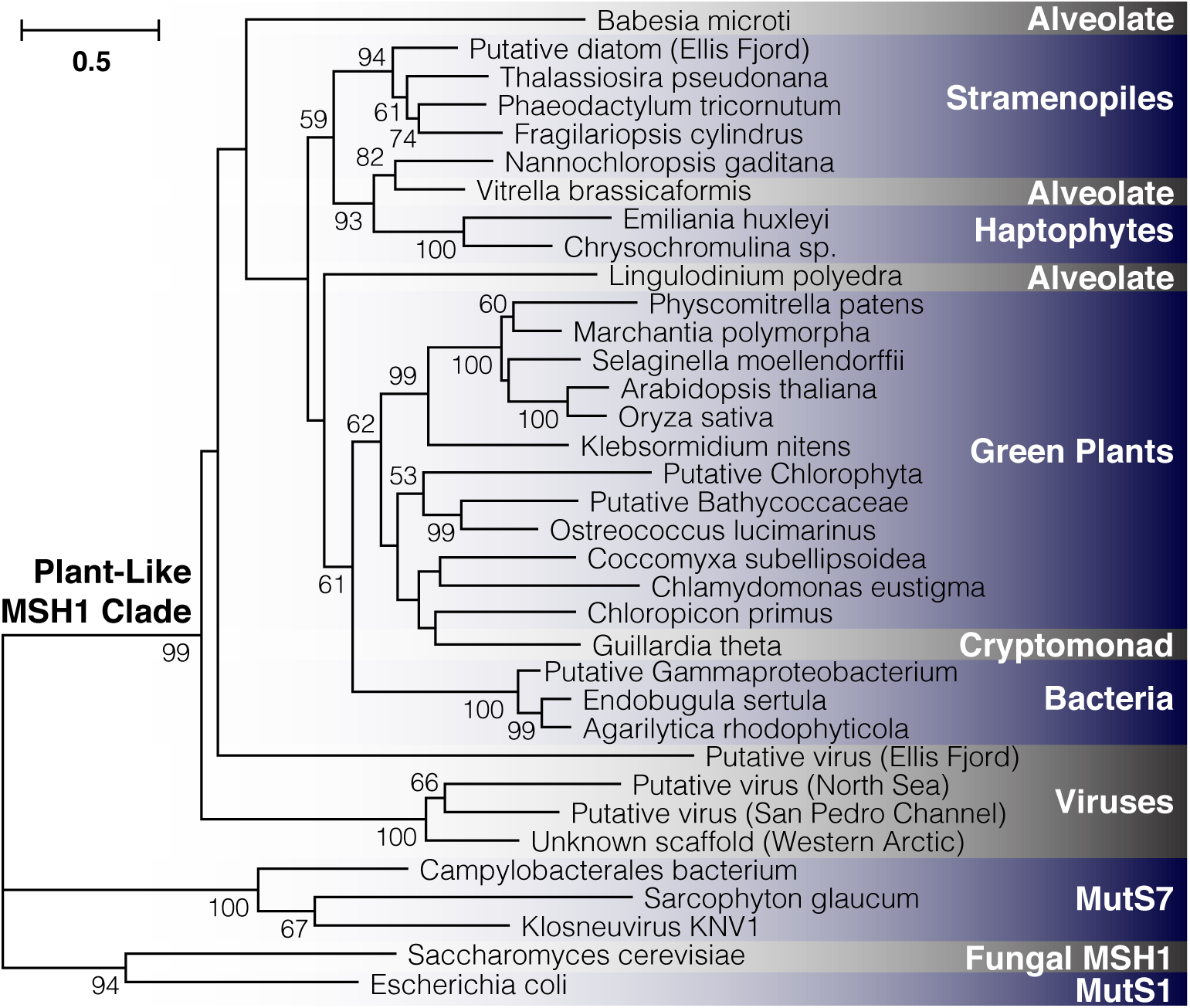
Detection of plant-like *MSH1* genes across diverse evolutionary lineages. The maximum-likelihood tree is constructed based on aligned protein sequences with branch lengths indicating the number of amino acid substitutions per site. Support values are percentages based on 1000 bootstrap pseudoreplicates (only values >50% are shown). Metagenomic samples are putatively classified based on other genes present on the same assembled contig (see Table S5 for information on sequence sources). The sample labeled “Unknown scaffold” lacks additional genes for classification purposes, but it clustered strongly with two sequences from the IMG/VR repository of viral genomic sequences.

## DISCUSSION

### The role of *MSH1* in maintaining the low mutation rates

The striking differences in mutation rates between cytoplasmic genomes in land plants vs. those in many other eukaryotes, including mammals, have posed a longstanding mystery because reactive oxygen species (ROS) are expected to be a potent source of DNA damage in all of these compartments. The presence of *MSH1* in plants and its dual targeting to the mitochondria and plastids may provide an explanation for their unusually low rates. Our findings that *msh1* mutants exhibit major increases in rates of cytoplasmic SNVs and indels align with a growing theme that the distinctive mutational properties of cytoplasmic genomes relative to the nucleus may be driven more by differences in RRR machinery than by ROS or the biochemical environment associated with cellular respiration and photosynthesis (44–46).

How does *MSH1* suppress cytoplasmic mutation rates? As a *mutS* homolog, it could conceivably be part of a conventional MMR pathway that has yet to be described in plant organelles. Postreplicative mismatch repair typically relies on the heuristic that mismatches in double-stranded DNA are more likely to reflect errors in the newly synthesized strand, with various mechanisms being used to specifically identify and repair that strand (47). However, the presence of a conventional MMR pathway would not explain how plant mitochondria and plastids maintain mutation rates substantially lower than in most eukaryotic genomes. An alternative, non-conventional pathway could involve use of the GIY-YIG endonuclease domain to introduce a DSB near sites identified by the mismatch recognition domain, followed by HDR of the DSB (9, 14, 16). This previously proposed model could lead to unusually high repair accuracy because it does not require use of a heuristic to determine which strand carries the error at mismatched sites, instead employing homologous recombination with an unaffected genome copy to “break the tie”.

A model based on DSBs and HDR might also explain some surprising features in our data. We found that the frequency of SNVs in wild type plants was much higher in the mitochondrial genome than the plastid genome, which is opposite the rates of evolution observed in these genomes on phylogenetic scales (1, 3). The mutation spectrum in wild type mitochondrial DNA was also dominated by GC→AT transitions (Fig. 1), which is inconsistent with the relatively neutral transition:transversion ratio observed in natural sequence variation both within and among species (24, 48, 49). We speculate that these apparent contrasts can be explained by the different copy numbers of mitochondrial and plastid genomes in vegetative tissues. Whereas, individual plastids each contain numerous genome copies, it has been estimated that there is less than one genome copy per mitochondrion in *Arabidopsis* leaf tissue (50). Therefore, even when *MSH1* is intact, HDR pathways may be less available for repair of mitochondrial DNA in vegetative tissues due to a paucity of homologous template copies (14), which would imply that the abundant GC→AT SNVs in wild type mitochondrial DNA are generally restricted to vegetative tissue and not transmitted to future generations. In contrast, the fusion of mitochondria into a large network within meristematic cells could provide an opportunity for mitochondrial genome copies to co-occur and utilize HDR (51). Because of the high genome copy number in plastids, they may rely more heavily on HDR even in vegetative tissue, which would explain why knocking out *MSH1* has a much larger proportional effect on observed variant frequencies in the plastid genome (Figs. 4 and S2). Therefore, we find growing support for the model in which *MSH1* is the link between mismatches, DSBs, and HDR. However, much remains to be done to validate this model, as researchers have yet to successfully express and purify full-length MSH1 for *in vitro* biochemical studies, and a recent analysis of the purified GIY-YIG domain was unable to detect endonuclease activity (30).

Which DNA aberrations does MSH1 recognize? The ability to bind to multiple types of disruptions in Watson-Crick pairing is a common feature of many MutS homologs (47). The fact that we observed increased frequencies of indels in *msh1* mutants implies that MSH1 can recognize indel loops, including in homopolymer regions, which are likely to be one of the most prevalent sources of polymerase errors, especially in the AT-rich genomes of plastids (52). The increased frequency of SNVs in *msh1* mutants also implies recognition of the bulges in DNA caused by mismatches and/or damaged bases. There is some evidence to suggest that MSH1 is capable of repairing both of these sources of mutation. The most prominent feature of the *msh1* mutation spectrum is the enormous increase in AT→GC transitions, which does not correspond to a major class of damage like cytosine deamination (GC→AT) or guanine oxidation (GC→TA). Therefore, this aspect of the mutation spectrum is more likely explained by polymerase misincorporations during DNA replication, although direct measurements of the spectrum of misincorporations by PolIA and PolIB would be needed to help test this hypothesis. We reasoned that disrupting the *POLIA* gene would increase mutation rates because of higher misincorporation rates for PolIB (9). The failure to find such an effect suggests a general insensitivity to polymerase errors when *MSH1* is intact, presumably because of its ability to recognize and repair these errors. This proposed role of *MSH1* would similarly explain why sequence evolution is so slow in these genomes despite polymerases with unusually high misincorporation rates (9). Disrupting genes involved in the repair of uracil (*UNG*) and 8-oxo-G (*FPG* and *OGG1*) failed to measurably affect the frequency of mitochondrial or plastid variants, which could indicate that MSH1 is capable of recognizing and correcting such damage. Alternatively, these sources of damage may be too minor under the tested growth conditions to contribute meaningfully to variant frequencies. But the fact that *MSH1* was recently shown to exhibit higher expression in *ung* mutants (14) points towards a capability to recognize damaged bases in addition to conventional mismatches.

### The evolutionary history of *MSH1* and parallels with other *mutS* lineages

To date, MSH1 has only been identified and studied in green plants. Researchers have previously noted the similarities in domain architecture between MSH1 and MutS7 (26), a lineage within the MutS family that independently acquired a C-terminal fusion of an endonuclease domain (39). MutS7 is encoded in the mitochondrial genome itself of octocorals, another eukaryotic lineage with unusually slow rates of mitochondrial genome evolution (53), as well as in the genomes of a small number of bacterial lineages and some giant viruses (54). In this sense, our results extend the parallels between MSH1 and MutS7 to include features of their phylogenetic distribution, as each are scattered across disparate lineages of eukaryotes, bacteria and viruses. The distribution of MSH1 (Fig. 7) clearly implies some history of horizontal gene transfer. However, the ancient divergences, sparse representation outside of eukaryotes, and poor phylogenetic resolution at deep splits within the gene tree make the timing of such events or specific donors and recipients unclear. Another open question is the functional role of MSH1 outside of land plants. The similarities in its effects on organelle genome stability between angiosperms and mosses (24, 28) suggest that much of the role of MSH1 in cytoplasmic genome maintenance are likely ancestral at least in land plants. Notably, all the eukaryotes that we identified as having MSH1 outside of green plants harbor a plastid derived from secondary endosymbiosis. It will therefore be interesting to assess whether it has mitochondrial and/or plastid functions in these eukaryotes (some show *in silico* targeting predictions to the organelles; Table S5). This pattern also raises the question as to whether it was ancestrally present deep in the eukaryotic tree and subsequently lost in many lineages or transferred among major eukaryotic lineages in conjunction with secondary endosymbiosis.

Because the apparent viral copies of *MSH1* were curated from metagenomic assemblies and bulk environmental virus sampling (Table S5), we were not able to assign these sequences to a specific type of virus. Interestingly, however, one of these cases was found on a viral-like metagenomic contig in the IMG/VR database that is >100 kb in size, and another co-occurs on a contig with a gene that has a top BLAST hit to the Mimiviridae, a clade of giant viruses. Therefore, similar to *mutS7*, it appears that *MSH1* may reside in giant viruses. We speculate that such viruses, which are also known as nucleocytoplasmic large DNA viruses or NCLDV (55), have acted as a repository for distinctive RRR machinery and a repeated source of horizontal acquisition by eukaryotic lineages, reshaping the mechanisms of cytoplasmic mutation rate and genome maintenance.

## MATERIALS AND METHODS

A complete description of the methods is available as supplementary material. In brief, mitochondrial and plastid DNA isolations were performed on rosette tissue harvested after seven to nine weeks of growth from either *A. thaliana* Col-0 or from F3 families derived from crossing mutant lines (Table S3) against *A. thaliana* Col-0 (Fig. 2). Duplex sequencing followed a modified version of the protocol of Kennedy et al. (37) that was first optimized in our lab by testing on single-colony *E. coli* samples. Sequencing performed on an Illumina NovaSeq 6000 at the University of Colorado Cancer Center. Data processing was performed with a custom pipeline available at https://github.com/dbsloan/duplexseq, and sequencing data are available via the NCBI Sequence Read Archive (PRJNA604834 and PRJNA604956). Inheritance of selected high-frequency SNVs in the F4 generation was assessed with droplet digital PCR on a Bio-Rad QX200 system, using fluorescently labeled allele-specific probes. To assess the phylogenetic distribution of plant-like *MSH1* genes, searches were performed against the NCBI nr protein database, as well as metagenomic and viral repositories hosted by JGI (42, 43). Identified sequences were used for maximum-likelihood phylogenetic analysis with PhyML v3.3.20190321.

## ACKNOWLEDGEMENTS

We thank Dolores Córdoba-Cañero for providing *fpg* and *ogg1* mutant *Arabidopsis* lines and Claudia Gentry-Weeks for providing the *E. coli* K12 MG1655 strain. We also thank Mychaela Hodous, Jocelyn Lapham, Holly Harroun, Amber Torres, and Mariella Rivera for assistance with plant growth, seed collection, and PCR genotyping. Jeff Palmer and members of the Sloan Lab also provided insightful comments and discussion. This work was supported by a grant from the National Institutes of Health (R01 GM118046) and a National Science Foundation NRT GAUSSI Graduate Fellowship (DGE-1450032).

## SUPPLEMENTARY TEXT

### Estimating noise threshold of duplex sequencing with *E. coli* single-colony analysis

Duplex sequencing has been used to detect rare variants by taking advantage of a reported error rate at or below 10^−7^ per bp (37). To assess the fidelity of this sequencing method in our hands, we used 2-ml liquid cultures each derived from a single colony of *Escherichia coli*. We chose these samples as our best approximation of a negative control that should be (nearly) free of true double-stranded mutations because of the low mutation rate in *E. coli* and the relatively small number of rounds of cell division required to reach saturation in a 2-ml volume (56). As such, we used these samples to estimate the duplex-sequencing error rate (while recognizing that this estimate may be conservatively high if some variants are true *de novo* mutations rather than sequencing errors). Because the standard method to fragment DNA samples with ultrasonication is known to introduce substantial oxidative damage (57, 58), we tested this approach alongside an alternative enzymatic fragmentation strategy (New England Biolabs dsDNA Fragmentase). We also performed each fragmentation strategy either with or without subsequent treatment with multiple DNA repair enzymes to eliminate common forms of single-stranded DNA damage. The unrepaired ultrasonication libraries showed the expected signature of oxidative damage dominated by single-stranded G→T errors (59), but much of these strand-specific effects could be effectively removed by enzymatic treatment (Fig. S7). For reasons that are unclear, Fragmentase treatment produced extremely high rates of single-stranded misincorporation of As (i.e., C→A, G→A, and T→A), as well as single-stranded indels, which were insensitive to subsequent repair treatment (Fig. S7). However, because these errors were generally not matched by a complementary change on the other strand, they were successfully filtered out during generation of DCS data. The average frequencies of SNVs in DCS data were statistically indistinguishable for repaired ultrasonication and Fragmentase libraries (Table S2). We used the ultrasonication methods with enzymatic repair for all subsequent experiments in this study. For this library type, the average frequency of SNVs across three *E. coli* biological replicates was 2.1 × 10^−8^ per bp, and there was only a single identified indel in a total of 240 Mb of mapped DCS data, confirming the extreme accuracy of duplex sequencing (Table S2).

## SUPPLEMENTARY MATERIALS AND METHODS

### *Arabidopsis* lines and growth conditions

*Arabidopsis thaliana* Col-0 was used as the wild type line used for all analyses. Existing mutant lines (Table S3) were used as pollen donors in crosses with Col-0 to generate heterozygous F1 individuals (Fig. 2). All mutant lines were originally generated in a Col-0 background with exception of the *fpg* and *ogg1* mutants, which were in a Landsberg *erecta* (L*er*) background (11, 60). F2 plants were screened with allele-specific PCR markers (Table S6) to identify individuals that were homozygous for the mutant allele and others that were homozygous for the wild type allele. Seeds were collected from each identified F2 homozygote to produce F3 full-sib families. After cold-stratification for three days, seeds were germinated and grown on ProMix BX soil mix in a growth room under a 10-hr short-day lighting conditions to extend rosette growth prior to bolting. For each candidate gene, sets of three F3 homozygous mutant families and three F3 wild type control families were grown in parallel. After seven to nine weeks of growth, approximately 35 g of rosette tissue was harvested from each family (representing approximately 60 individuals per family) and used for mitochondrial and plastid DNA purification. Additional F3 individuals in each family were left unharvested and allowed to set seed for subsequent analysis of F4 individuals.

### Mitochondrial and plastid DNA isolation

Mitochondrial DNA purification was performed as described previously (49) except that initial mitochondrial pelleting spins were done at 20,000 rcf and subsequent washing spins were performed at 25,800 rcf. Plastid DNA was isolated simultaneously from the same tissue sample, using interleaved centrifugation steps during the mitochondrial DNA extraction. Pellets containing plastids (chloroplasts) from the 1500 rcf spin in the mitochondrial extraction protocol were gently resuspended with a paintbrush in a total of 6 ml of wash buffer (0.35 M sorbitol, 50 mM Tris-HCl pH 8, 25 mM EDTA) and then loaded onto two discontinuous sucrose gradients with 9 ml of 30% sucrose solution on top of 19.5 ml of 52% sucrose solution, each containing 50 mM Tris-HCl pH 8 and 25 mM EDTA. The sucrose gradients were then centrifuged at 95,400 rcf for 1.5 hr in a JS24.38 swinging bucket rotor on a Beckman-Coulter Avanti JXN-30 centrifuge. Plastids were harvested from the interface between the 30% and 52% sucrose solutions, diluted in wash buffer, and centrifuged in a JA14.50 fixed-angle rotor at 25,800 rcf for 16 min. Plastids were washed two more times by gently resuspending pellets in wash buffer using a paintbrush and centrifuging at 25,800 rcf. All organelle isolation steps were performed at 4° C in a cold room or refrigerated centrifuge. Plastid lysis and DNA purification followed the same protocol as described previously for mitochondrial samples (49). Tissues samples were processed in pairs, with one mutant and one wild type sample in each batch.

### *E. coli* growth and DNA extraction

A glycerol stock of the *E. coli* K12 MG1655 strain was streaked onto an LB (Luria-Bertani) agar plate and grown overnight at 37° C. Single colonies were then used to inoculate each of three 2-ml liquid LB cultures, which were grown overnight at 37 °C on a shaker at 200 rpm. Half of each culture (approximately 4 × 10^9^ cells) was used for DNA extraction with an Invitrogen PureLink Genomic DNA Kit, following the manufacturer’s protocol for gram-negative bacteria.

### Duplex sequencing library construction and Illumina sequencing

Master stocks of duplex sequencing adapters were generated following the specific quantities and protocol described by Kennedy et al. (37) with the oligos in Table S7. For each mitochondrial and plastid DNA sample, a total of 100 ng was diluted in 50 μl of T_10_E_0.1_ buffer (10 mM Tris-HCl pH 8, 0.1 mM EDTA). DNA was fragmented with the Covaris M220 Focused-Ultrasonicator in microTUBE AFA Fiber Screw-Cap tubes to a target size of approximately 250 bp, using a duty cycle of 20%, peak incident power of 50 W, 200 cycles/burst, and six bouts of shearing for 20 sec each (separated by 15 sec pauses) at a temperature of 6° C. Ultrasonication settings were adapted from the protocol of Schmitt et al. (61).

After fragmentation, 80 ng of DNA was end repaired with the NEBNext End Repair Module (New England Biolabs E6050S) for 30 min at 20° C. Samples were then cleaned with 1.6 volumes of solid phase reversible immobilization (SPRI) beads. A-tailing of eluted samples was performed in a 50 μl reaction volume, containing 5 U Klenow Fragment enzyme (New Biolabs M0212), 1 mM dATP, and 1x NEB Buffer 2, at 37°C for 1 hr, followed by clean-up with 1.6 volumes of SPRI beads. After quantification with a Qubit dsDNA HS Kit (Invitrogen), adapter ligation was performed on 20 ng of the resulting sample in a 50 μl reaction volume, using the NEBNext Quick Ligation Module (New England Biolabs E6056S) and 1 μl of a 64-fold dilution of the duplex sequencing adapter master stock, for 15 min at 20° C. Following adapter ligation, samples were cleaned with 0.8 volumes of SPRI beads. Half of the cleaned sample was then treated with a cocktail of repair enzymes to remove single-stranded damage in a 50 μl reaction volume containing 1x NEB CutSmart Buffer, 8 U Fpg (New England Biolabs M0240), 5 U Uracil-DNA Glycosylase (New England Biolabs M0280), and 10 U Endonuclease III (New England Biolabs M0268) for 30 min at 37° C. Samples were then cleaned with 1.6 volumes of SPRI beads.

The repaired product was quantified with a Qubit dsDNA HS Kit, and 50 pg was amplified and dual-indexed using the primers shown in Table S7 and the NEBNext Ultra II Q5 Master Mix (New England Biolabs M0544) according to the manufacturer’s instructions. All libraries were amplified for 19 cycles, which yielded the necessary redundancy to generate DCS families. After amplification, libraries were processed with 1 volume of SPRI beads and eluted in 20 μl T_10_E_0.1_ buffer. Libraries were assessed with an Agilent TapeStation 2200 and High Sensitivity D1000 reagents. If adapter dimers were detected (which was only the case for the batch of 12 libraries for *polIb* mutants and matched wild type controls), they were size selected with a 2% gel on a BluePippin (Sage Science), using a specified target range of 300-700 bp.

In initial tests of duplex sequencing with *E. coli* DNA sample, the above protocol was applied either with the described repair enzyme treatment or a control treatment that did not include these enzymes. It was also performed either with ultrasonication-based fragmentation or with an alternative fragmentation protocol based on dsDNA Fragmentase (New England Biolabs M0348). These protocol variations were performed in a 2×2 factorial design with three biological replicates. For the Fragmentase approach, 400 ng of DNA was incubated for 20 min at 37° C. Fragmentation was terminated by adding 5 μl of 0.5 M EDTA to each reaction. Samples were then cleaned with 1.6 volumes of SPRI beads.

Libraries were sequenced on 2×150 bp runs on an Illumina NovaSeq 6000 platform, with a target yield of 40M read pairs per library. Libraries were generally processed and sequenced in batches of 12 corresponding to each candidate gene and its matched wild type controls (2 genomes × 2 genotypes × 3 biological replicates). The only exception was the *msh1*-CS3246 set, for which the mitochondrial and plastid libraries were generated and sequenced in separate batches of six libraries. Library construction and sequencing of the original *A. thaliana* Col-0 families and the *E. coli* samples were also done in their own batches. The raw Illumina sequencing reads have been deposited to the NCBI Sequence Read Archive under BioProjects PRJNA604834 (*E. coli*) and PRJNA604956 (*Arabidopsis*). Individual accessions for each library are provided in Tables S1 and S4.

### Duplex sequencing data analysis

Raw Illumina reads from duplex sequencing libraries were processed with a custom Perl-based pipeline available at https://github.com/dbsloan/duplexseq. In the first step in the pipeline, 3′ read trimming for low quality bases (q20) and adapter sequence was performed with cutadapt v1.16 (62). The minimum length for retaining reads after trimming was set to 75, and the error tolerance for adapter trimming was set to 0.15. BBMerge (63) was then used to join overlapping paired-end reads into a single sequence where possible, with a minimum overlap of 30 bp and a maximum of five mismatches. The random duplex sequencing tags were then extracted from the resulting trimmed and merged reads, applying a stringent filter that rejected any reads with a barcode that contained a base with a quality score below 20. Reads were also filtered if they lacked the expected TGACT linker sequence built into the duplex sequencing adapters. Reads were then collapsed into single-stranded consensus sequences (SSCS), requiring a minimum of three reads to form an SSCS family. To call a consensus base, a minimum of 80% agreement was required within an SSCS family. When complementary SSCS families were available (reflecting the two different strands of an original double-stranded DNA molecule), they were used to form a DCS family. Any disagreements between the two complementary SSCS families were left as ambiguities in the DCS read, and any DCS read with ambiguities was later filtered out from downstream analyses.

The filtered DCS data were mapped using bowtie2 v2.2.3 under default parameter settings. The *E. coli* data were mapped against the corresponding K12 MG1655 reference genome (GenBank U00096.3). We later had to exclude a called SNV at position 4,296,060 and indel at position 4,296,381 because they were shared across all three replicates and appeared to reflect fixed differences between our *E. coli* line and the reference. Likewise, in *Arabidopsis*, we found that the plastid genome in our Col-0 line had a 1-bp expansion in a homopolymer at position 28,673 relative to the published reference genome (GenBank NC_000932.1). In this case, we updated the reference genome for mapping purposes such that all reported coordinates for variants reflect a 1-bp shift at that position. For the mitochondrial genome reference, we used our recently revised version (64) of the Col-0 sequence (GenBank NC_037304.1). All *Arabidopsis* samples were mapped to a database that contained both the mitochondrial and plastid genomes to avoid cross-mapping due to related sequences shared between them because of historical intergenomic transfers (MTPTs). The resulting mapping (SAM) files were parsed to extract all SNVs and simple indels, as well as coverage data. Multi-nucleotide variants (MNVs) and more complicated structural variants were not analyzed in this pipeline.

The identified variants and associated coverage data were filtered to address known sources of errors and artefacts. First, because of the DNA fragmentation and end-repair steps involved in library construction, the positions near the ends of inserts can be prone to sequencing errors that falsely appear to be true double-stranded changes (37). Therefore, we excluded any variants and sequencing coverage associated with 10 bp at each end of a DCS read. Second, contaminating sequences can easily be mistaken for *de novo* mutations. In the case of mitochondrial and plastid genomes, one of the most likely sources of contamination is NUMT and NUPT sequences in the nucleus (38). Therefore, we used NCBI BLASTN v2.2.29+ to map each DCS reads containing an identified variant against the TAIR10 release of the *A. thaliana* Col-0 nuclear genome. Any variants that returned a perfect match to the nucleus were excluded as presumed NUMTs or NUPTs. However, one additional challenge for the mitochondrial genome is that *A. thaliana* Col-0 harbors a recent genome-scale insertion of mitochondrial DNA into Chromosome 2, only a fraction of which is accurately captured in the published nuclear genome assembly (65). Therefore, some NUMT artefacts cannot be detected using the currently available reference nuclear genome. To address this problem, we used total-cellular shotgun DNA sequencing data from *A. thaliana* Col-0 that was generated in a different lab (NCBI SRA SRR5216995) and therefore unlikely to share inherited heteroplasmic variants with our Col-0 line. We used these raw reads to generate a database of *k*-mer counts (*k* = 39 bp) with KMC v3.0.0 (66), and we checked all identified variants for presence in this database. We filtered all variants with a count of 30 or greater in the SRR5216995 dataset, as we found that this was a reliable threshold for distinguishing known NUMTs from background sequencing errors in the mitochondrial genome. Finally, an additional complication with identifying *de novo* SNVs and indels in the mitochondrial genome is that it contains an abundance of small to medium-sized repeats that can become recombinationally active in some of the mutants analyzed in this study (29, 34) and can exhibit rare recombination even in wild type genotypes (67). When chimeric sequences resulting from recombination are mapped against a reference genome, they can give the false indication that *de novo* point mutations or indels have occurred. To eliminate these false positives, we used NCBI BLASTN to map DCS reads containing identified variants against the reference mitochondrial genome with a maximum e-value of 1e-10 to check for secondary hits (i.e., related repeat sequences) that contained the exact variant and thus could have arisen by recombination between repeat copies in the genome. SNVs that met these criteria were removed from the variant call set as likely recombinants. In the case of indels, subsequent manual curation was required for all flagged candidates to confirm variants that were consistent with recombination because of the inconsistent handling of gaps in repetitive regions by BLAST.

Identified SNVs were further characterized based on the reference genome sequence and annotation to classify their location as protein-coding, rRNA, tRNA, intronic, or intergenic. For protein-coding variants, the effect (if any) on amino acid sequence was reported. We also extracted the flanking 5′ and 3′ bases to assess the effect of trinucleotide context on the occurrence of mutations.

The above steps were automated with the aforementioned tools available at https://github.com/dbsloan/duplexseq. The duplexseq_batchscripts.pl script in that repository can generate a shell script for each input library for submission to standard Slurm-based queuing systems with the same parameter settings that we applied for initial read processing and raw variant calling. Scripts are also provided to aggregate and filter the raw variant files from multiple samples run in parallel. Variant frequencies were calculated from output data by dividing the total number reads with an identified variant type by the relevant DCS coverage (expressed in total bp). Reported statistical analyses were performed in R v3.4.3, and plots were generated with the ggplot2 package.

### Analysis of mitochondrial and plastid genome coverage variation in *Arabidopsis* mutants

To investigate region-specific changes in copy number in mitochondrial and plastid genomes for *Arabidopsis* mutants relative to wild type, we used the DCS reads generated above to calculate sequence coverage in terms of counts per million mapped reads as described previously (49). Counts were averaged over 500-bp windows, and means were taken across three biological replicates for both mutants and matched wild type controls. Thus, when the reported ratio of these values (Figs. 3, S1 and S3) exceeds a value of 1, it indicates a region with increased relative coverage within the genome in the mutant compared to wild type. Likewise, values below 1 indicate decreased relative coverage in mutants.

### Expression and intron splicing analysis for *msh1-*SALK_046763 mutants

To test the hypothesis that *msh1* SALK_046763 mutants exhibited weaker effects on leaf variegation and mutation rates because their intronic T-DNA insert only reduces but does eliminate *MSH1* expression, we sampled F4 individuals derived from F3 homozygous SALK_046763 mutant families and from their matched F3 wild type controls (Fig. 2). Four F4 individuals were sampled from each genotype (including at least one from all three F3 families for both mutants and wild types). Approximately 60-90 mg of rosette leaf tissue was collected from each plant after approximately 8 weeks of growth under 10-hr short-day lighting conditions, flash frozen with liquid nitrogen, and immediately processed using the Qiagen RNeasy Plant Mini Kit with on-column DNase digestion. For each sample, 1 μg of RNA was reverse transcribed into cDNA using Bio-Rad iScript Reverse Transcription Supermix in a 20 μl reaction volume. Quantitative PCR (qPCR) was performed with two different *MSH1* markers – one spanning the exons that flank the T-DNA insertion in intron 8 and another in exon 16 – as well as two reference gene markers (Table S8). All primer pairs were tested with conventional endpoint PCR and gel electrophoresis to ensure amplification of a single product of expected size. Primer pair efficiency was assessed using a dilution series (Table S8). qPCR reactions (20 μl total volume) contained 10 μl of Bio-Rad 2x iTaq SYBR Green Supermix, 10 pmole of each primer, and 1 μl of cDNA. Reactions were run on the Bio-Rad CFX96 Touch Real-Time PCR System. Thermal cycling conditions included 95° C for 3 min followed by 40 cycles of 95° C for 10 sec and 60° C for 30 sec, with a final melt curve analysis ramping from 65-95° C. Three technical replicates were run for each of the four biological replicates. In addition, a single replicate of a no-reverse-transcriptase control was run for each plant sample, and one no-template control was run for each primer set. For each cDNA sample, an average threshold cycle (C_T_) value was calculated from the three technical reps. The geometric mean of the two reference genes was calculated to create a single reference C_T_ value for each of the eight plants for normalization (calculation of ΔC_T_ values). Differences in *MSH1* expression between mutant and wild type genotypes were estimated using the ΔΔC_T_ method.

To test whether properly spliced *MSH1* mRNA transcripts were produced in SALK_046763 homozygous mutants, we performed Sanger sequencing of an RT-PCR product spanning the junction of exons 8 and 9. cDNA was generated as described above for qPCR experiments. Endpoint PCR was performed using NEBNext 2x Master Mix, 0.25 μM of forward and reverse primers (MSH1 Exon 5F: 5′–CTGGTCTCAATCCTTTTGGTG–3′ and MSH1 Exon 10R: 5′– CAAACTCTCCCCAGCGGC–3′) and 1 μl of cDNA template in a 20 μl reaction volume. cDNAs were amplified from all four sampled SALK_046763 homozygous mutant plants. Thermal cycling conditions were as follows: 98° C for 30 sec; 35 cycles of 98° C for 10 sec, 60° C for 15 sec and 72° C for 20 sec; 72° C for 2 min. PCR products were visualized by gel electrophoresis to ensure the amplification of a single product of the expect size (~460 bp). For Sanger sequencing reactions, 2.5 μl of PCR product was treated with 1 μl of ExoSAP-IT (Thermo-Fisher) and incubated at 37° C for 15 min after which the enzymes were deactivated at 80° C for 15 min. Each treated PCR sample was sent to GeneWiz for Sanger sequencing after addition of 5 μl of MSH1 Exon 5F primer and 6.5 μl of dH_2_0. A single representative electropherogram for the junction between exons 8 and 9 is shown in Fig. S6, but all samples confirmed the presence of properly spliced products.

### ddPCR heteroplasmy assays

To assess the possibility that observed heteroplasmies in duplex sequencing data could be transmitted across generations, we grew F4 seed collected from eight individuals from each of the three *msh1*-CS3246 mutant F3 families used in duplex sequencing. These F3 parents were siblings of the actual F3 individuals that were harvested for the duplex sequencing analysis. Approximately 80 mg of rosette leaf tissue was collected from F4 plants after approximately 8 weeks of growth under 10-hr short-day lighting conditions. Collected tissue was either immediately processed or stored at −80° C until processing. Tissue samples were disrupted using the Qiagen TissueLyser, and total-cellular DNA was extracted using the Qiagen Plant DNeasy Mini Kit. DNA was quantified using a Qubit dsDNA HS Kit.

Locus-specific primers (Table S9) and allele-specific fluorescently labeled probes (Table S10) were designed for five different SNV targets (one mitochondrial and four plastid), which represented five of the most abundant variants in the *msh1*-CS3246 mutant lines based on duplex sequencing read counts (Dataset S1). Probes were designed with the target SNV in the center.

Primers were tested by conventional endpoint PCR and gel electrophoresis to ensure that a single band was amplified for each primer set. Each ddPCR reaction was set up in an initial 20 μl volume composed of 1x Bio-Rad ddPCR Supermix for Probes (no dUTP), 250 nM final concentration of each probe, 900 nM final concentration of each primer, 1 μl of the restriction enzyme BglII (Thermo Scientific FD0083), and 5 µl of diluted template DNA (5-500 pg depending on the SNV target and type of DNA, i.e., total cellular or organellar). The restriction enzyme was included for fragmentation of genomic DNA to improve ddPCR efficiency and was selected because it does not cut within any of the target amplicons. PCR emulsions were created with a Bio-Rad QX200 Droplet Generator according to manufacturer’s instructions, using Bio-Rad DG8 Cartridges and QX200 Droplet Generation Oil for Probes. Amplification was performed in a Bio-Rad C1000 Touch Thermal Cycler with a deep-well block with the following program: enzyme activation at 95° C for 10 min, 40 cycles of 94° C for 30 sec and a variable annealing/extension temperature (see Table S9) for 1 min, and enzyme deactivation at 98° C for 10 min – with a ramp speed of 2° C per sec for all steps. Droplets were read on the Bio-Rad QX200 Droplet Reader and analyzed using QuantaSoft Analysis Software to calculate copy numbers of reference and alternative alleles in each sample. Channel thresholds were set based on initial experiments utilizing positive and negative controls.

### MSH1 phylogenetic analysis

To assess the distribution of *MSH1* outside of green plants, we performed BLASTP searches with the *Arabidopsis* MSH1 protein sequence against the NCBI nr database using taxonomic filters to exclude Viridiplantae. We also used individual searches restricted to specific clades, including Bacteria, Archaea, Glaucophyta, Rhodophyta, and Opisthokonta. Candidates for plant-like MSH1 proteins were identified based on high amino-acid identity and near full-length hits that extended through the characteristic GIY-YIG domain. To further expand our search to include some of the vast amount of biological diversity that is unculturable and only detected in environmental samples, we queried a sample of 2000 metagenome assemblies from the JGI IMG/MER repository (43). We also searched against the IMG/VR database, which houses the largest available collection of viral sequences from both sequenced isolates and environmental samples (42). In cases where MSH1*-*like sequences were identified on metagenomic scaffolds, we searched other proteins encoded in the flanking sequence against the NCBI nr database to infer possible origins for the scaffold.

Identified protein sequences were aligned with other select members of the MutS family (Table S5) using the E-INS-i algorithm in MAFFT v6.903b (68). The resulting alignments were trimmed with Gblocks v0.91b (http://molevol.cmima.csic.es/castresana/Gblocks.html) to remove low-quality alignment regions, using the following parameters: t=p; b1=18; b2=18; b3=10; b4=5; b5=h. Models of sequence evolution were assessed with ProtTest v3.4.2 (69), which identified LG+I+G+F as the preferred model based on the Akaike Information Criterion. A maximum-likelihood phylogenetic search was then performed in PhyML v3.3.20190321 (70) using this substitution model, an SPR search of tree space, 1000 random starts, and 1000 bootstrap replicates.

---

**Figure S1.**
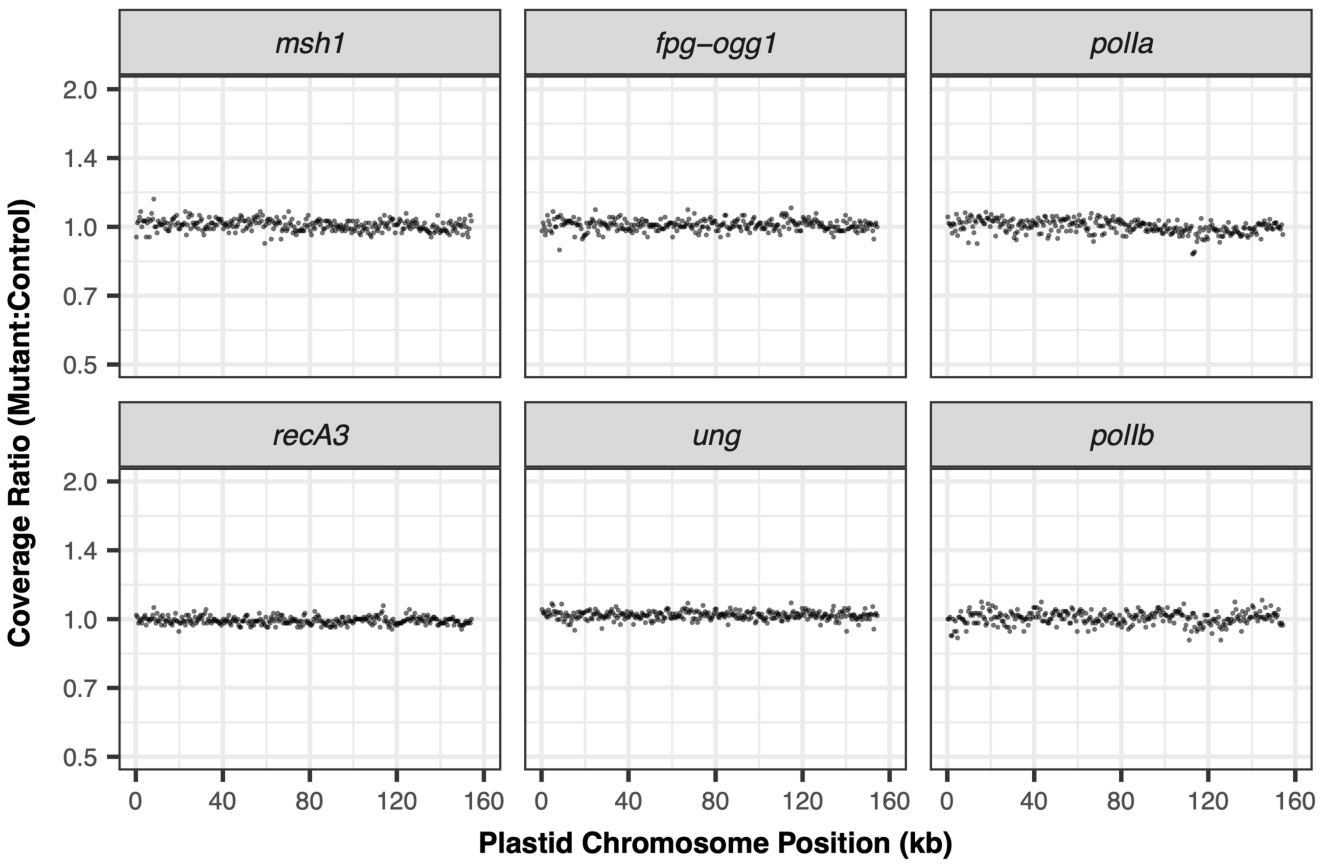
Sequencing coverage variation across the plastid genome in mutants relative to their matched wild type controls. Each panel represents an average of three biological replicates, with the exception of two cases where a single outlier replicate (*ung* mutant 3 and *POLIA* wild type 3) was excluded due to what appeared to be unusually high amplification bias. The reported ratios are based on counts per million mapped reads in 500-bp windows.

**Figure S2.**
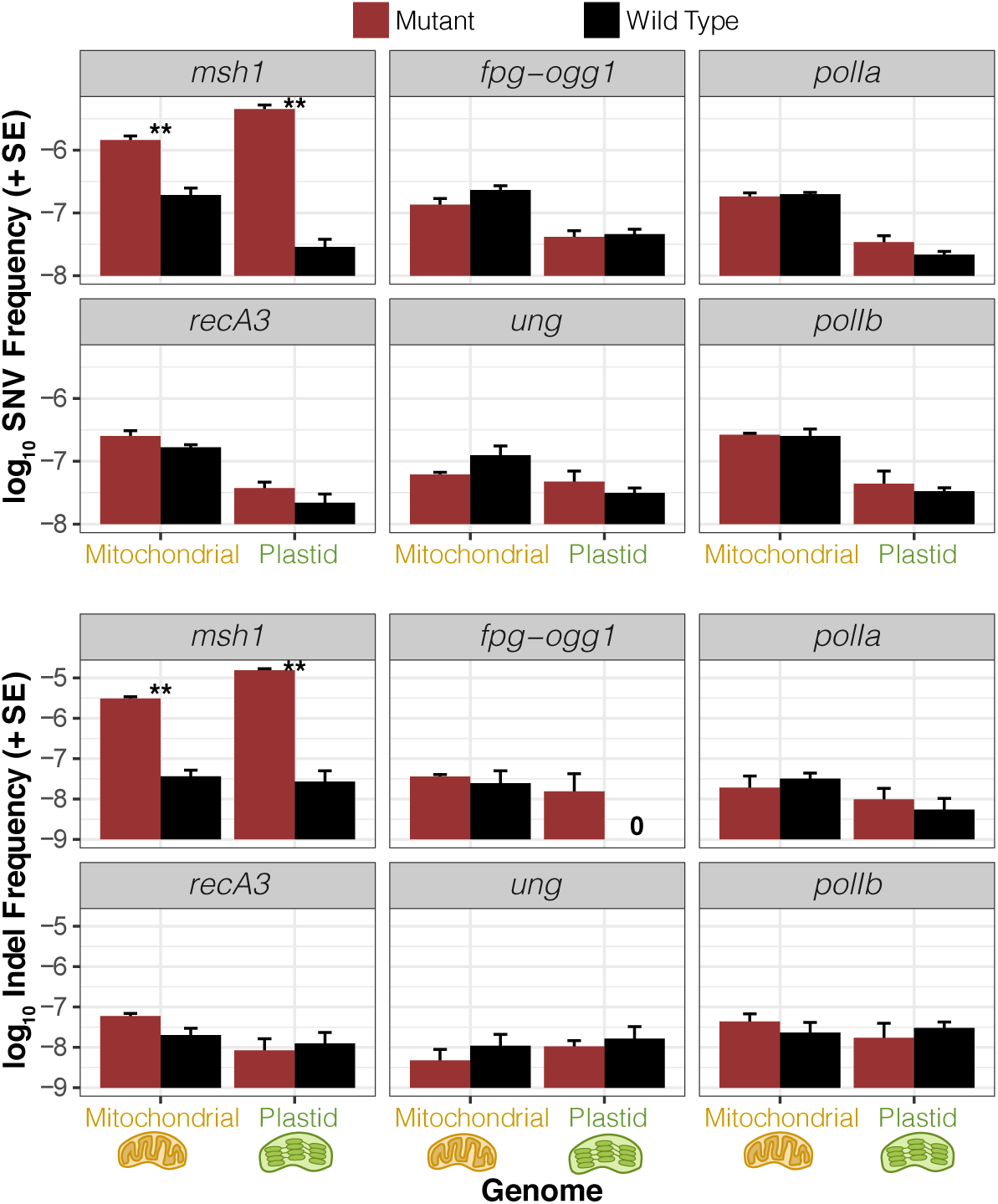
The same data on SNV and indel frequencies presented in Fig. 4 but plotted on a log scale. See Fig. 4 legend for additional information.

**Figure S3.**
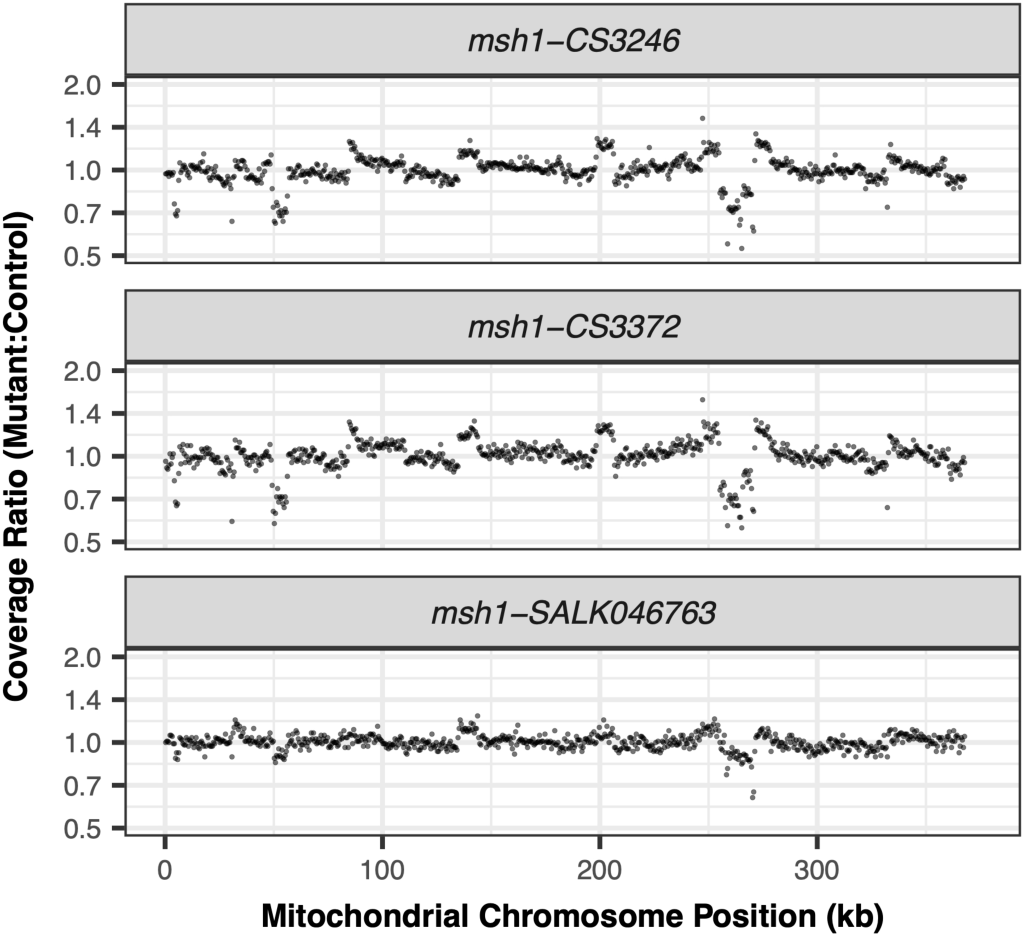
Sequencing coverage variation across the mitochondrial genome in three different *msh1* mutants relative to their matched wild type controls. Each panel represents an average of three biological replicates. The reported ratios are based on counts per million mapped reads in 500-bp windows. The weaker effects of SALK_046763 likely reflect the fact that this allele has a reduced expression level but is not a full functional knockout.

**Figure S4.**
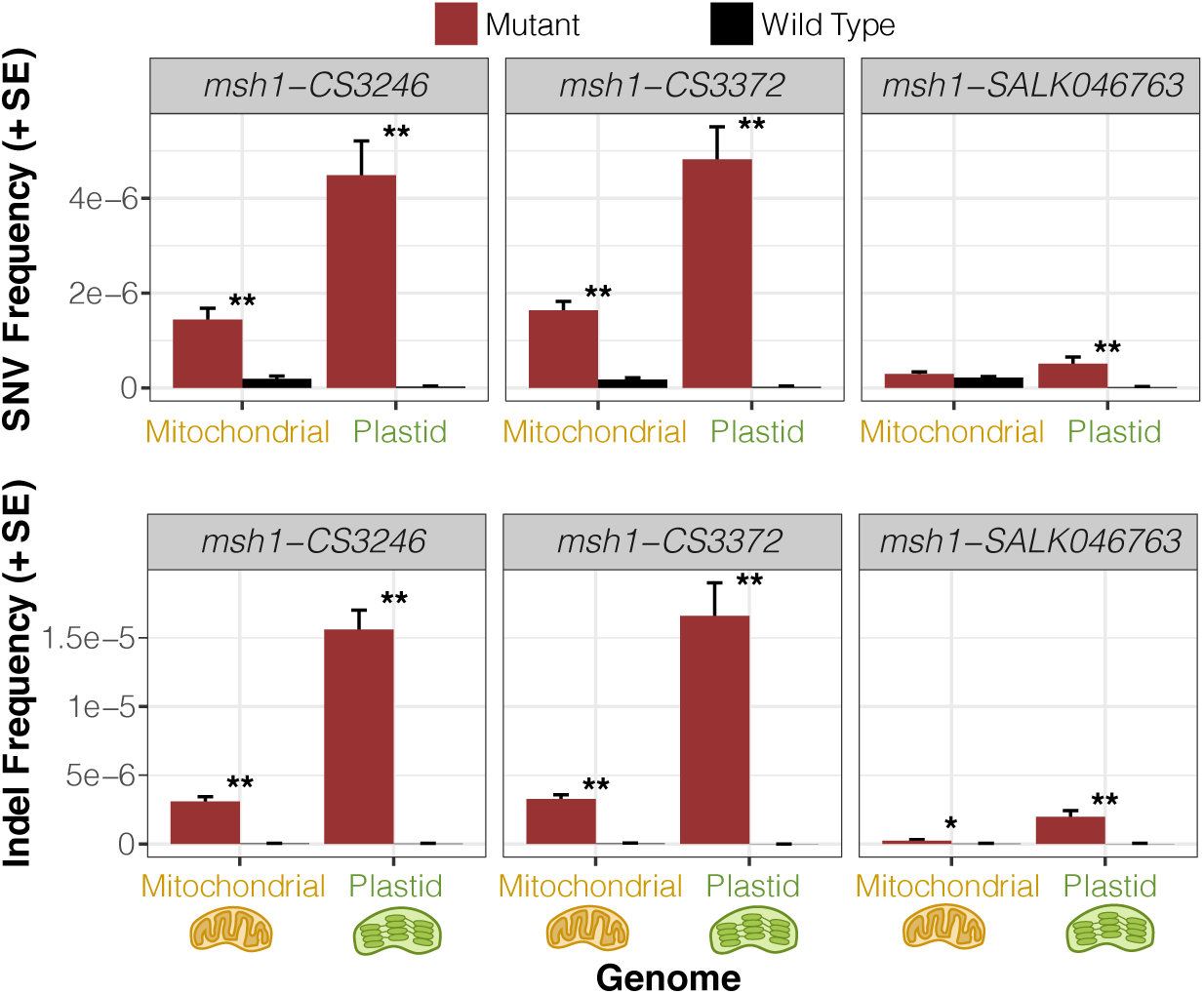
Observed frequency of mitochondrial and plastid SNVs (top) and indels (bottom) based on duplex sequencing in three different *Arabidopsis msh1* mutant backgrounds compared to matched wild type controls. Variant frequencies are calculated as the total number of observed mismatches or indels in mapped duplex consensus sequences divided by the total bp of sequence coverage. Means and standard errors are based on three replicate F3 families for each genotype (see Fig. 2). Significant differences between mutant and wild type genotypes at a level of *P* < 0.05 or *P* < 0.01 (t-tests on log-transformed values) are indicated by * and **, respectively. The weaker effects of SALK_046763 likely reflect the fact that this allele has a reduced expression level but is not a full functional knockout.

**Figure S5.**
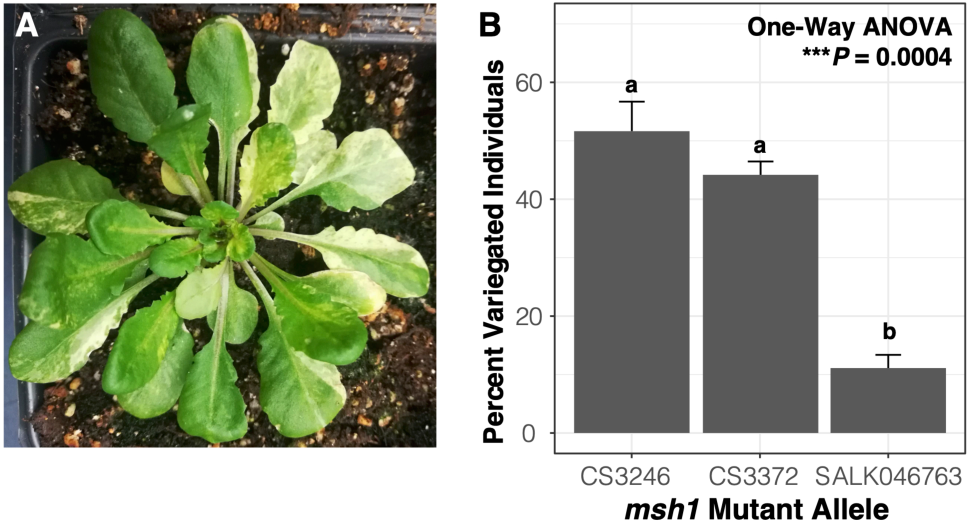
Extent of leaf variegation observed for different *msh1* mutant alleles. **A.** An example of an *msh1* mutant (CS3372) individual with a leaf-variegation phenotype. **B.** Values represent the percent of individuals in an F3 family from a homozygous mutant F2 parent that showed visible leaf variegation at time of harvest for mitochondrial and plastid DNA extraction. Means and standard errors are from three replicate F3 families from each mutant line (see Fig. 2). Between 45 and 66 individuals were scored for each family. Lowercase letters indicate significant differences between alleles based on a Tukey’s HSD test. Consistent with its lower rate of observed sequence and structural variation in cytoplasmic genomes, the SALK_046763 *msh1* mutant line exhibited less severe phenotypic effects.

**Figure S6.**
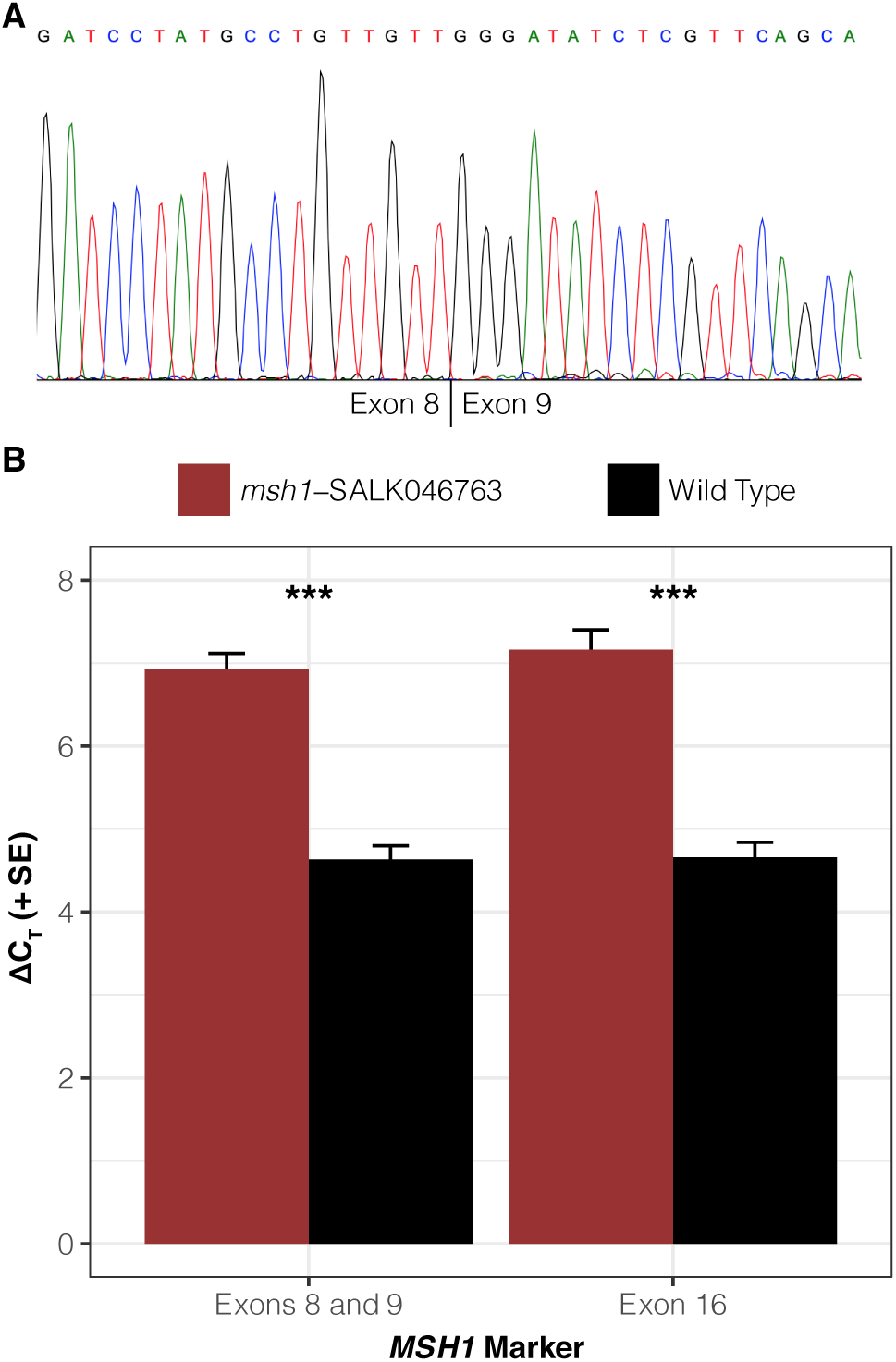
Intact *MSH1* transcripts but reduced expression level in homozygous SALK_046763 *msh1* mutants. **A.** Sanger trace from cDNA sequencing confirms that properly spliced transcripts are present in SALK_046763 *msh1* mutants despite the large T-DNA insertion in intron 8. The vertical line below the trace indicates the location of the expected splice junction between exons 8 and 9. **B.** ΔC_T_ values are calculated based on the difference in quantitative reverse-transcriptase PCR (qRT-PCR) threshold cycle value for each indicated *MSH1* marker and the geometric mean of the threshold cycle values from two reference genes (*UBC* and *UBC9*). Means and standard errors are from four biological replicates (F4 plants derived from crossing design described in Fig. 2), each of which is based on the mean of three technical replicates. The SALK_046763 mutants exhibit higher ΔC_T_ (indicating lower *MSH1* expression). Both *MSH1* markers indicate a similar shift in ΔC_T_ values (2.3 cycles for exons 8/9 and 2.5 cycles for exon 16), corresponding to an approximately 5-fold difference in transcript abundance. Significant differences between mutant and wild type genotypes at a level of *P* < 0.001 (*t*-tests) are indicated by ***.

**Figure S7.**
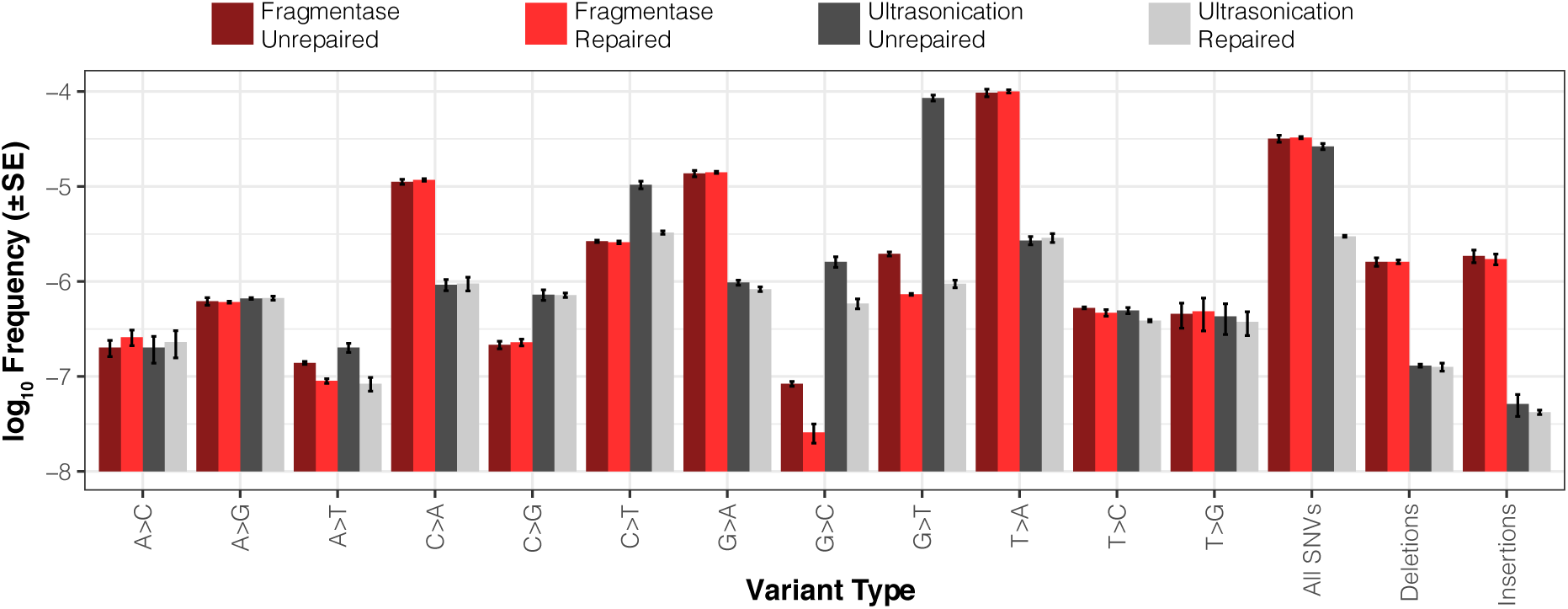
Strand-specific variant frequencies from *E. coli* SSCS (not DCS) data for different library construction preparation methods (shearing by dsDNA Fragmentase or Covaris ultrasonication, either with or without subsequent enzymatic repair treatment). Values are based on means and standard errors from three biological replicates.

**Table S1.** *E. coli* duplex sequencing library summary

| Library (SRA) | Frag. Method | Repair Treatment | Sample | Read Pairs (M) | Raw Sequence (Gb) | Mapped DCS (Mb) |
| --- | --- | --- | --- | --- | --- | --- |
| SRR11018568 | Fragmentase | Yes | 1 | 30.7 | 9.2 | 74.8 |
| SRR11018564 | Fragmentase | Yes | 2 | 35.9 | 10.8 | 79.1 |
| SRR11018562 | Fragmentase | Yes | 3 | 27.0 | 8.1 | 75.6 |
| SRR11018567 | Fragmentase | No | 1 | 36.4 | 10.9 | 95.9 |
| SRR11018563 | Fragmentase | No | 2 | 39.6 | 11.9 | 97.8 |
| SRR11018561 | Fragmentase | No | 3 | 28.7 | 8.6 | 86.2 |
| SRR11018560 | Ultrasonication | Yes | 1 | 37.0 | 11.1 | 80.2 |
| SRR11018558 | Ultrasonication | Yes | 2 | 27.2 | 8.1 | 68.9 |
| SRR11018566 | Ultrasonication | Yes | 3 | 28.7 | 8.6 | 90.9 |
| SRR11018559 | Ultrasonication | No | 1 | 32.6 | 9.8 | 98.6 |
| SRR11018557 | Ultrasonication | No | 2 | 21.4 | 6.4 | 69.9 |
| SRR11018565 | Ultrasonication | No | 3 | 28.9 | 8.7 | 65.5 |
| **Total** |  |  |  | **374.1** | **112.2** | **983.2** |

**Table S2.** Variants detected in *E. coli* duplex consensus sequence data for different library construction preparation methods (shearing by dsDNA Fragmentase or Covaris ultrasonication, either with or without enzymatic repair treatment). Three different DNA samples were collected and then subjected to each of the methods.

| Method | Sample | Coverage (bp) | AT>CG | AT>GC | AT>TA | GC>AT | GC>CG | GC>TA | All SNVs | SNV Freq | Indels |
| --- | --- | --- | --- | --- | --- | --- | --- | --- | --- | --- | --- |
| Fragmentase<br>Unrepaired | A | 9.6E+07 | 0 | 0 | 0 | 0 | 0 | 0 | 0 | 0 | 0 |
|  | B | 9.8E+07 | 0 | 0 | 0 | 0 | 0 | 1 | 1 | 1.0E-08 | 1 |
|  | C | 8.6E+07 | 0 | 0 | 0 | 1 | 0 | 1 | 2 | 2.3E-08 | 0 |
|  | Total | 2.8E+08 | 0 | 0 | 0 | 1 | 0 | 2 | 3 | 1.1E-08 | 1 |
| Fragmentase<br>Repaired | A | 7.5E+07 | 0 | 0 | 0 | 1 | 0 | 0 | 1 | 1.3E-08 | 0 |
|  | B | 7.9E+07 | 1 | 0 | 0 | 0 | 0 | 0 | 1 | 1.3E-08 | 0 |
|  | C | 7.6E+07 | 0 | 0 | 0 | 0 | 0 | 0 | 0 | 0 | 0 |
|  | Total | 2.3E+08 | 1 | 0 | 0 | 1 | 0 | 0 | 2 | 8.7E-09 | 0 |
| Ultrasonication<br>Unrepaired | A | 9.9E+07 | 1 | 1 | 0 | 0 | 2 | 2 | 6 | 6.1E-08 | 0 |
|  | B | 7.0E+07 | 0 | 0 | 0 | 0 | 2 | 2 | 4 | 5.7E-08 | 0 |
|  | C | 6.5E+07 | 0 | 0 | 0 | 0 | 4 | 5 | 9 | 1.4E-07 | 0 |
|  | Total | 2.3E+08 | 1 | 1 | 0 | 0 | 8 | 9 | 19 | 8.1E-08 | 0 |
| Ultrasonication<br>Repaired | A | 8.0E+07 | 0 | 0 | 0 | 0 | 3 | 0 | 3 | 3.7E-08 | 0 |
|  | B | 6.9E+07 | 0 | 1 | 0 | 0 | 0 | 0 | 1 | 1.5E-08 | 0 |
|  | C | 9.1E+07 | 0 | 0 | 0 | 1 | 0 | 0 | 1 | 1.1E-08 | 1 |
|  | Total | 2.4E+08 | 0 | 1 | 0 | 1 | 3 | 0 | 5 | 2.1E-08 | 1 |

**Table S3.**
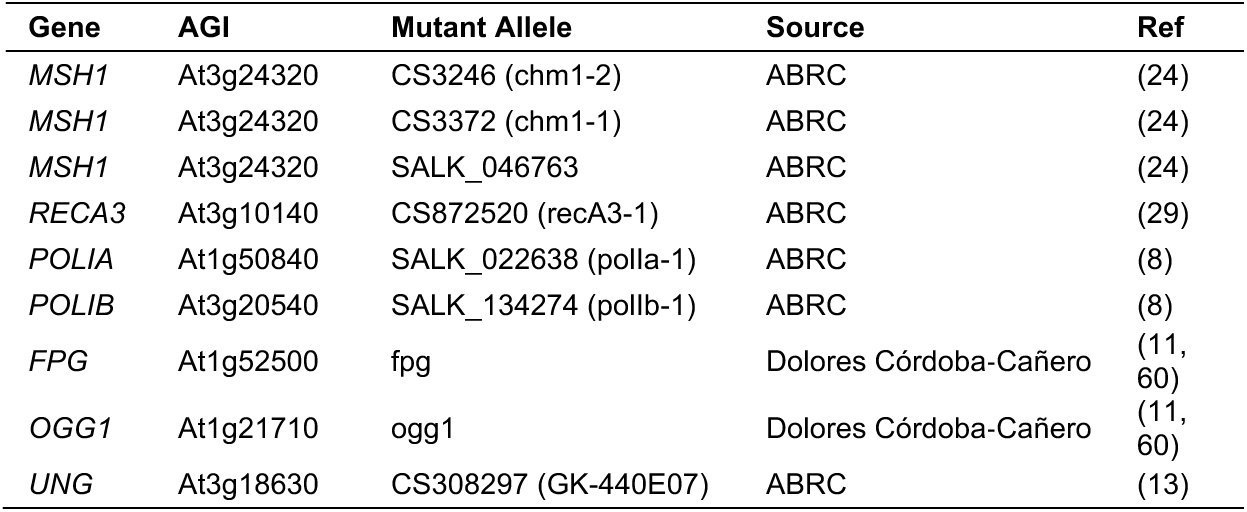
Mutant lines used for analysis of candidate genes involved in DNA replication, recombination, and repair.

**Table S4.** *Arabidopsis* duplex sequencing library summary

| Library (SRA) | Gene/Line | Genotype | Purification | Rep | Read Pairs (M) | Raw Seq (Gb) | Mapped DCS (Mb) |  |
| --- | --- | --- | --- | --- | --- | --- | --- | --- |
|  |  |  |  |  |  |  | Mito | Plastid |
| SRR11025108 | Col-0 | . | cpDNA | 1 | 26.1 | 7.8 | 5.4 | 54.6 |
| SRR11025107 | Col-0 | . | cpDNA | 2 | 22.9 | 6.9 | 3.4 | 41.3 |
| SRR11025106 | Col-0 | . | cpDNA | 3 | 21.8 | 6.6 | 10.3 | 61.4 |
| SRR11025105 | Col-0 | . | mtDNA | 1 | 23.6 | 7.1 | 39.3 | 3.6 |
| SRR11025175 | Col-0 | . | mtDNA | 2 | 25.8 | 7.7 | 33.7 | 4.8 |
| SRR11025174 | Col-0 | . | mtDNA | 3 | 22.1 | 6.6 | 30.8 | 2.1 |
| SRR11025121 | fpg-ogg1 | Mutant | cpDNA | 1 | 29.8 | 8.9 | 6.4 | 53.4 |
| SRR11025120 | fpg-ogg1 | Mutant | cpDNA | 2 | 23.9 | 7.2 | 3.9 | 35.3 |
| SRR11025119 | fpg-ogg1 | Mutant | cpDNA | 3 | 30.5 | 9.2 | 5.7 | 61.9 |
| SRR11025118 | fpg-ogg1 | Mutant | mtDNA | 1 | 22.7 | 6.8 | 42.8 | 1.0 |
| SRR11025117 | fpg-ogg1 | Mutant | mtDNA | 2 | 33.7 | 10.1 | 58.9 | 2.8 |
| SRR11025116 | fpg-ogg1 | Mutant | mtDNA | 3 | 30.9 | 9.3 | 50.4 | 3.0 |
| SRR11025114 | fpg-ogg1 | Wild Type | cpDNA | 1 | 39.4 | 11.8 | 11.8 | 74.3 |
| SRR11025113 | fpg-ogg1 | Wild Type | cpDNA | 2 | 33.2 | 10.0 | 7.0 | 56.6 |
| SRR11025112 | fpg-ogg1 | Wild Type | cpDNA | 3 | 27.6 | 8.3 | 6.0 | 44.9 |
| SRR11025111 | fpg-ogg1 | Wild Type | mtDNA | 1 | 33.7 | 10.1 | 57.8 | 1.6 |
| SRR11025110 | fpg-ogg1 | Wild Type | mtDNA | 2 | 32.0 | 9.6 | 32.0 | 1.6 |
| SRR11025109 | fpg-ogg1 | Wild Type | mtDNA | 3 | 32.7 | 9.8 | 39.8 | 2.4 |
| SRR11025100 | msh1-CS3246 | Mutant | cpDNA | 1 | 42.4 | 12.7 | 28.7 | 60.8 |
| SRR11025099 | msh1-CS3246 | Mutant | cpDNA | 2 | 32.4 | 9.7 | 25.4 | 71.5 |
| SRR11025098 | msh1-CS3246 | Mutant | cpDNA | 3 | 43.9 | 13.2 | 25.9 | 106.7 |
| SRR11025093 | msh1-CS3246 | Mutant | mtDNA | 1 | 32.6 | 9.8 | 103.1 | 2.7 |
| SRR11025092 | msh1-CS3246 | Mutant | mtDNA | 2 | 44.5 | 13.3 | 125.4 | 4.6 |
| SRR11025091 | msh1-CS3246 | Mutant | mtDNA | 3 | 41.5 | 12.5 | 98.8 | 2.4 |
| SRR11025097 | msh1-CS3246 | Wild Type | cpDNA | 1 | 41.3 | 12.4 | 30.0 | 53.6 |
| SRR11025095 | msh1-CS3246 | Wild Type | cpDNA | 2 | 38.1 | 11.4 | 30.3 | 73.3 |
| SRR11025094 | msh1-CS3246 | Wild Type | cpDNA | 3 | 35.5 | 10.6 | 18.7 | 98.0 |
| SRR11025090 | msh1-CS3246 | Wild Type | mtDNA | 1 | 38.9 | 11.7 | 126.6 | 1.5 |
| SRR11025089 | msh1-CS3246 | Wild Type | mtDNA | 2 | 40.2 | 12.1 | 117.9 | 2.2 |
| SRR11025088 | msh1-CS3246 | Wild Type | mtDNA | 3 | 36.3 | 10.9 | 143.7 | 1.4 |
| SRR11025159 | msh1-CS3372 | Mutant | cpDNA | 1 | 40.2 | 12.1 | 12.4 | 98.4 |
| SRR11025158 | msh1-CS3372 | Mutant | cpDNA | 2 | 39.8 | 11.9 | 12.9 | 95.7 |
| SRR11025157 | msh1-CS3372 | Mutant | cpDNA | 3 | 36.1 | 10.8 | 17.3 | 77.4 |
| SRR11025156 | msh1-CS3372 | Mutant | mtDNA | 1 | 33.5 | 10.1 | 79.8 | 8.3 |
| SRR11025155 | msh1-CS3372 | Mutant | mtDNA | 2 | 32.8 | 9.8 | 64.4 | 3.2 |
| SRR11025154 | msh1-CS3372 | Mutant | mtDNA | 3 | 36.6 | 11.0 | 62.0 | 16.2 |
| SRR11025152 | msh1-CS3372 | Wild Type | cpDNA | 1 | 38.6 | 11.6 | 6.2 | 79.2 |
| SRR11025151 | msh1-CS3372 | Wild Type | cpDNA | 2 | 33.6 | 10.1 | 7.7 | 76.6 |
| SRR11025104 | msh1-CS3372 | Wild Type | cpDNA | 3 | 38.4 | 11.5 | 11.8 | 90.5 |
| SRR11025103 | msh1-CS3372 | Wild Type | mtDNA | 1 | 35.6 | 10.7 | 56.2 | 2.4 |
| SRR11025102 | msh1-CS3372 | Wild Type | mtDNA | 2 | 44.7 | 13.4 | 78.1 | 1.8 |
| SRR11025101 | msh1-CS3372 | Wild Type | mtDNA | 3 | 32.0 | 9.6 | 48.7 | 15.9 |
| SRR11025087 | msh1-SALK046763 | Mutant | cpDNA | 1 | 29.1 | 8.7 | 5.2 | 24.8 |
| SRR11025086 | msh1-SALK046763 | Mutant | cpDNA | 2 | 26.2 | 7.9 | 14.5 | 39.8 |
| SRR11025078 | msh1-SALK046763 | Mutant | cpDNA | 3 | 36.2 | 10.9 | 17.5 | 64.0 |
| SRR11025083 | msh1-SALK046763 | Mutant | mtDNA | 1 | 30.1 | 9.0 | 61.2 | 1.0 |
| SRR11025082 | msh1-SALK046763 | Mutant | mtDNA | 2 | 34.9 | 10.5 | 70.6 | 1.2 |
| SRR11025149 | msh1-SALK046763 | Mutant | mtDNA | 3 | 29.9 | 9.0 | 82.4 | 1.7 |
| SRR11025080 | msh1-SALK046763 | Wild Type | cpDNA | 1 | 35.0 | 10.5 | 14.1 | 55.8 |
| SRR11025079 | msh1-SALK046763 | Wild Type | cpDNA | 2 | 32.8 | 9.8 | 13.9 | 63.1 |
| SRR11025084 | msh1-SALK046763 | Wild Type | cpDNA | 3 | 27.6 | 8.3 | 9.4 | 28.3 |
| SRR11025077 | msh1-SALK046763 | Wild Type | mtDNA | 1 | 33.8 | 10.1 | 84.7 | 2.2 |
| SRR11025150 | msh1-SALK046763 | Wild Type | mtDNA | 2 | 30.7 | 9.2 | 62.8 | 1.1 |
| SRR11025081 | msh1-SALK046763 | Wild Type | mtDNA | 3 | 27.9 | 8.4 | 59.3 | 1.0 |
| SRR11025172 | polla | Mutant | cpDNA | 1 | 32.0 | 9.6 | 4.3 | 60.8 |
| SRR11025171 | polla | Mutant | cpDNA | 2 | 42.1 | 12.6 | 13.7 | 121.1 |
| SRR11025170 | polla | Mutant | cpDNA | 3 | 47.5 | 14.2 | 7.5 | 130.6 |
| SRR11025169 | polla | Mutant | mtDNA | 1 | 35.6 | 10.7 | 45.9 | 7.4 |
| SRR11025168 | polla | Mutant | mtDNA | 2 | 55.6 | 16.7 | 97.9 | 14.3 |
| SRR11025167 | polla | Mutant | mtDNA | 3 | 40.6 | 12.2 | 84.2 | 4.4 |
| SRR11025166 | polla | Wild Type | cpDNA | 1 | 39.1 | 11.7 | 6.0 | 96.8 |
| SRR11025165 | polla | Wild Type | cpDNA | 2 | 47.6 | 14.3 | 9.6 | 135.0 |
| SRR11025163 | polla | Wild Type | cpDNA | 3 | 41.9 | 12.6 | 14.4 | 105.2 |
| SRR11025162 | polla | Wild Type | mtDNA | 1 | 54.1 | 16.2 | 60.4 | 7.1 |
| SRR11025161 | polla | Wild Type | mtDNA | 2 | 35.4 | 10.6 | 61.8 | 13.3 |
| SRR11025160 | polla | Wild Type | mtDNA | 3 | 50.2 | 15.1 | 115.3 | 3.1 |
| SRR11025178 | pollb | Mutant | cpDNA | 1 | 23.1 | 6.9 | 11.1 | 31.3 |
| SRR11025177 | pollb | Mutant | cpDNA | 2 | 40.1 | 12.0 | 27.7 | 103.7 |
| SRR11025164 | pollb | Mutant | cpDNA | 3 | 23.7 | 7.1 | 14.5 | 22.5 |
| SRR11025153 | pollb | Mutant | mtDNA | 1 | 38.5 | 11.6 | 117.6 | 2.2 |
| SRR11025096 | pollb | Mutant | mtDNA | 2 | 39.3 | 11.8 | 87.3 | 1.7 |
| SRR11025085 | pollb | Mutant | mtDNA | 3 | 44.6 | 13.4 | 102.8 | 1.0 |
| SRR11025148 | pollb | Wild Type | cpDNA | 1 | 21.9 | 6.6 | 14.6 | 28.1 |
| SRR11025137 | pollb | Wild Type | cpDNA | 2 | 50.6 | 15.2 | 37.2 | 114.9 |
| SRR11025126 | pollb | Wild Type | cpDNA | 3 | 24.2 | 7.3 | 15.4 | 22.8 |
| SRR11025115 | pollb | Wild Type | mtDNA | 1 | 38.9 | 11.7 | 143.4 | 2.4 |
| SRR11025176 | pollb | Wild Type | mtDNA | 2 | 43.5 | 13.1 | 105.5 | 1.6 |
| SRR11025173 | pollb | Wild Type | mtDNA | 3 | 52.1 | 15.6 | 123.9 | 1.4 |
| SRR11025147 | recA3 | Mutant | cpDNA | 1 | 62.7 | 18.8 | 26.3 | 197.4 |
| SRR11025146 | recA3 | Mutant | cpDNA | 2 | 55.0 | 16.5 | 21.2 | 171.4 |
| SRR11025145 | recA3 | Mutant | cpDNA | 3 | 57.3 | 17.2 | 21.7 | 179.7 |
| SRR11025144 | recA3 | Mutant | mtDNA | 1 | 56.7 | 17.0 | 127.1 | 5.9 |
| SRR11025143 | recA3 | Mutant | mtDNA | 2 | 53.6 | 16.1 | 144.7 | 5.3 |
| SRR11025142 | recA3 | Mutant | mtDNA | 3 | 53.7 | 16.1 | 131.5 | 13.6 |
| SRR11025141 | recA3 | Wild Type | cpDNA | 1 | 55.2 | 16.6 | 26.5 | 175.2 |
| SRR11025140 | recA3 | Wild Type | cpDNA | 2 | 51.4 | 15.4 | 19.1 | 162.3 |
| SRR11025139 | recA3 | Wild Type | cpDNA | 3 | 64.1 | 19.2 | 34.0 | 184.7 |
| SRR11025138 | recA3 | Wild Type | mtDNA | 1 | 56.8 | 17.0 | 167.2 | 4.8 |
| SRR11025136 | recA3 | Wild Type | mtDNA | 2 | 60.7 | 18.2 | 175.2 | 4.7 |
| SRR11025135 | recA3 | Wild Type | mtDNA | 3 | 47.3 | 14.2 | 170.8 | 7.2 |
| SRR11025134 | ung | Mutant | cpDNA | 1 | 69.3 | 20.8 | 28.5 | 129.0 |
| SRR11025133 | ung | Mutant | cpDNA | 2 | 62.5 | 18.8 | 21.8 | 121.3 |
| SRR11025132 | ung | Mutant | cpDNA | 3 | 55.9 | 16.8 | 16.0 | 109.1 |
| SRR11025131 | ung | Mutant | mtDNA | 1 | 56.4 | 16.9 | 124.7 | 2.6 |
| SRR11025130 | ung | Mutant | mtDNA | 2 | 57.3 | 17.2 | 106.8 | 14.2 |
| SRR11025129 | ung | Mutant | mtDNA | 3 | 56.3 | 16.9 | 103.1 | 2.1 |
| SRR11025128 | ung | Wild Type | cpDNA | 1 | 63.2 | 19.0 | 26.3 | 122.5 |
| SRR11025127 | ung | Wild Type | cpDNA | 2 | 61.4 | 18.4 | 19.3 | 113.1 |
| SRR11025125 | ung | Wild Type | cpDNA | 3 | 56.3 | 16.9 | 11.2 | 91.2 |
| SRR11025124 | ung | Wild Type | mtDNA | 1 | 47.2 | 14.1 | 125.5 | 2.3 |
| SRR11025123 | ung | Wild Type | mtDNA | 2 | 45.8 | 13.8 | 73.2 | 2.3 |
| SRR11025122 | ung | Wild Type | mtDNA | 3 | 73.2 | 22.0 | 142.7 | 10.7 |
| **Total** |  |  |  |  | **4137.3** | **1241.2** | **5459.2** | **4700.7** |

**Table S5.** MSH1 and other MutS sequences used for phylogenetic analysis

| Sequence | Clade | Source | Accession | TargetP Prediction |
| --- | --- | --- | --- | --- |
| *Arabidopsis thaliana* | Viridiplantae | GenBank | AAO49798 | Mitochondrial |
| *Oryza sativa* | Viridiplantae | GenBank | XP_015636674 | Plastid |
| *Selaginella moellendorffii* | Viridiplantae | GenBank | XP_024525391 | Mitochondrial |
| *Physcomitrella patens* | Viridiplantae | GenBank | XP_024362883 | Mitochondrial |
| *Marchantia polymorpha* | Viridiplantae | GenBank | PTQ35957 | None |
| *Klebsormidium nitens* | Viridiplantae | GenBank | GAQ89439 | None |
| *Chlamydomonas eustigma* | Viridiplantae | GenBank | GAX83118 | Plastid |
| *Chloropicon primus* | Viridiplantae | GenBank | QDZ22437 | None |
| *Coccomyxa subellipsoidea* C-169 | Viridiplantae | GenBank | XP_005644580 | None |
| *Ostreococcus lucimarinus* | Viridiplantae | GenBank | XP_001416776 | None |
| Putative Bathycoccaceae (Tara Oceans metagenome) | Viridiplantae | JGI IMG/MER | Ga0211588_10010061 | None |
| Putative Chlorophyta (Trout Bog Lake metagenome) | Viridiplantae | JGI IMG/MER | Ga0164297_100118835 | None |
| *Guillardia theta* | Cryptophyta | GenBank | XP_005824881 | None |
| *Chrysochromulina tobinii* | Haptophyta | GenBank | KOO23129 | Mitochondrial |
| *Emiliania huxleyi* CCMP 1516 | Haptophyta | JGI IMG | 2508101639 | Mitochondrial |
| *Babesia microti* | Alveolata | GenBank | XP_021338715 | Mitochondrial |
| *Lingulodinium polyedra* | Alveolata | GenBank | QDO16335 | None |
| *Vitrella brassicaformis* | Alveolata | GenBank | CEL93038 | None |
| Putative diatom (Ellis Fjord metagenome) | Stramenopile | IMG/MER | Ga0307978_10152622 | None |
| *Thalassiosira pseudonana* | Stramenopile | GenBank | XP_002294874 | None |
| *Fragilariopsis cylindrus* | Stramenopile | GenBank | OEU09203 | None |
| *Phaeodactylum tricornutum* | Stramenopile | GenBank | XP_002177023 | None |
| *Nannochloropsis gaditana* | Stramenopile | GenBank | XP_005855452 | Mitochondrial |
| *Agarilytica rhodophyticola* | Gammaproteobacteria | GenBank | WP_086930438 | None |
| *Endobugula sertula* | Gammaproteobacteria | GenBank | ODS23082/ODS23083 | None |
| Putative bacterium (Damariscotta River mineral coupon metagenome) | Gammaproteobacteria | JGI IMG/MER | Ga0316202_100001462 | . |
| Putative virus (North Sea metagenome) | Virus | JGI IMG/VR | Ga0115564_1000006915 | . |
| Putative virus (San Pedro Channel metagenome) | Virus | JGI IMG/VR | Ga0181430_100039610 | . |
| Putative virus (Ellis Fjord metagenome) | Virus | JGI IMG/MER | Ga0307978_10013766 | . |
| Unknown Scaffold (Western Arctic Ocean metagenome) | Undetermined | JGI IMG/MER | Ga0302121_100079582 | . |
| *Sarcophyton glaucum* | MutS7 | GenBank | O63852 | . |
| *Klosneuvirus KNV1* | MutS7 | GenBank | ARF11760 | . |
| *Campylobacteriales* bacterium 16-40-21 | MutS7 | GenBank | OYZ35361 | . |
| *Saccharomyces cerevisiae* | Fungal MSH1 | GenBank | AJU17961 | Mitochondrial |
| *Escherichia coli* | MutS1 | GenBank | AIZ90229 | . |

**Table S6.**
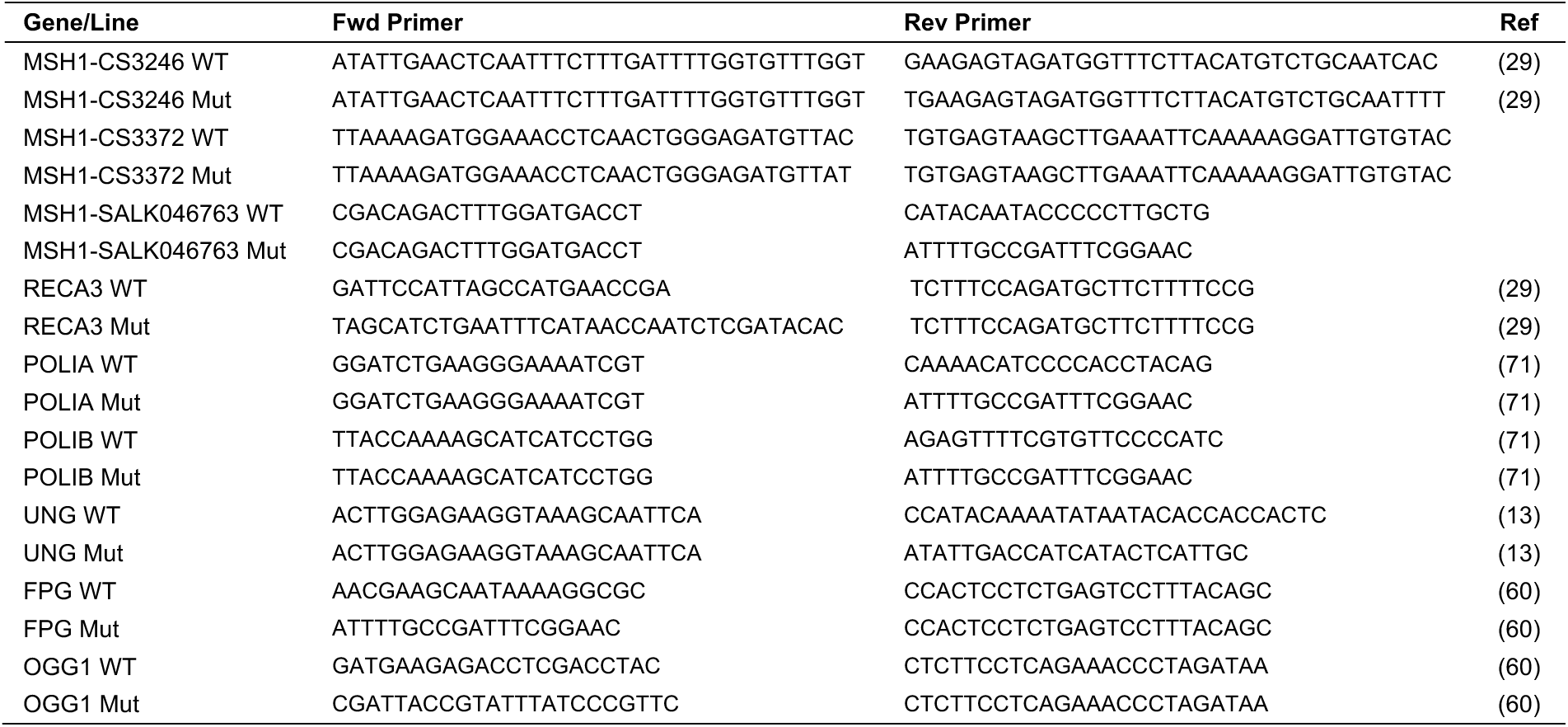
Primers for genotyping of mutant alleles for candidate genes.

**Table S7.**
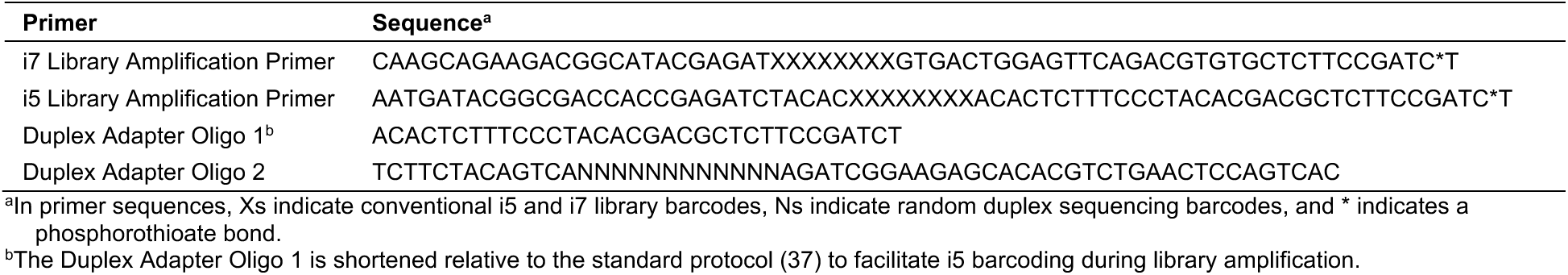
Oligos for duplex sequencing adapters and library amplification

**Table S8.**
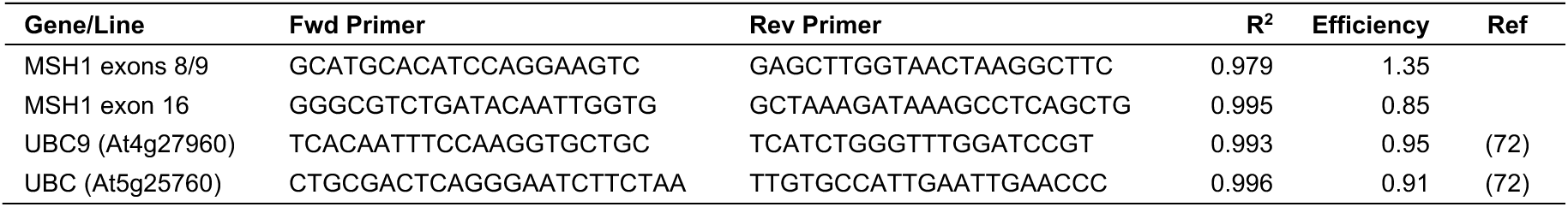
Primers for *msh1*-SALK046763 qPCR expression analysis and efficiency statistics from dilution series

**Table S9.**
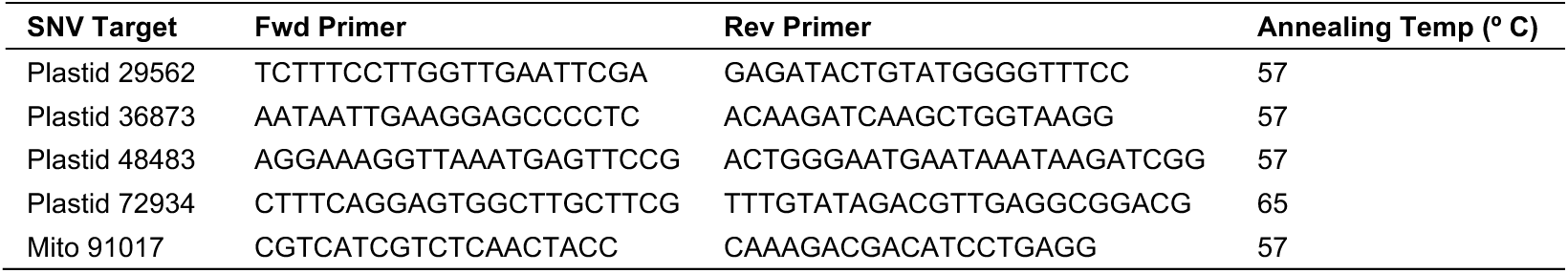
Primers for ddPCR heteroplasmy assays

**Table S10.** Allele-specific probes for ddPCR heteroplasmy assays (synthesized by Integrated DNA Technologies [IDT])

| SNV Target | Reference Probe<sup>a*</sup> | Alternative Probe<sup>b*</sup> |
| --- | --- | --- |
| Plastid 29562 | /56-FAM/CCTATT+CC+A+T+TTT+C+CT/3IABkFQ/ | /5HEX/CC+T+ATT+CC+A+C+TTTCC/3IABkFQ/ |
| Plastid 36873 | /56-FAM/C+CAT+T+CT+G+T+CTAAATAG/3IABkFQ/ | /5HEX/C+CATTCT+G+C+CTA+A+ATA/3IABkFQ/ |
| Plastid 48483 | /56-FAM/TT+GCC+C+T+TCAA+C+TAT/3IABkFQ/ | /5HEX/TTGCC+C+C+TCAAC+TAT/3IABkFQ/ |
| Plastid 72934 | /56-FAM/CAA+A+ACC+C+T+CCACG+CC/3IABkFQ/ | /5HEX/CAAA+ACC+C+C+CCACGC/3IABkFQ/ |
| Mito 91017 | /56-FAM/TCAACTAG+A+A+TT+C+C+CTT/3IABkFQ/ | /5HEX/CAA+CTAG+A+G+TTC+CCT/3IABkFQ/ |

<sup>a</sup>Each probe for the reference allele carries a 5' 6-FAM fluorescent modification and a 3' Iowa Black fluorescent quencher.
<sup>b</sup>Each probe for the alternative allele carries a 5' HEX fluorescent modification and a 3' Iowa Black fluorescent quencher.
\*Bases preceded by a symbol + indicate locked nucleic acids (LNA).

**Dataset S1.** Detailed summary information for each variant call. Coordinates are based on the *A. thaliana* Col-0 mitochondrial reference genome (NC_037304.1) and a modified version of the plastid reference genome (NC_000932.1) that includes a 1-bp insertion at position 28,673. (DatasetS1.xlsx)

